# Molecular evolution of toothed whale genes reveals adaptations to echolocating in different environments

**DOI:** 10.1101/2023.01.10.523466

**Authors:** L. Magpali, E. Ramos, A. Picorelli, L. Freitas, M.F. Nery

## Abstract

**Background:** Echolocation was a key development in toothed whale evolution, enabling their adaptation and diversification across various environments. Previous bioacoustic and morphological studies suggest that environmental pressures have influenced the evolution of echolocation in toothed whales. This hypothesis demands further investigation, especially regarding the molecular mechanisms involved in the adaptive radiation of toothed whales across multiple habitats. Here we show that the coding sequences of four hearing genes involved in echolocation (*CDH23*, *SLC26A5*, *TMC1,* and *CLDN14*) have different signatures of molecular evolution among riverine, coastal, and oceanic dolphins, suggesting that the evolutionary constraints of these habitats shaped the underlying genetic diversity of the toothed whale sonar.

**Results:** Our comparative analysis across 37 odontocete species revealed patterns of accelerated evolution within coastal and riverine lineages, supporting the hypothesis that shallow habitats pose specific selective pressures to sonar propagation, which are not found in the deep ocean. All toothed whales with genes evolving under positive selection are shallow coastal species, including three species that have recently diverged from freshwater lineages (*Cephalorhynchus commersonii*, *Sotalia guianensi*s, and *Orcaella heinsohni* - *CDH23*), and three species that operate specialized Narrow Band High Frequency (NBHF) Sonars (*Phocoena sinus* - *SLC26A5*, *Neophocaena phocaenoides* and *Cephalorhynchus commersonii* - *CDH23*). For river dolphins and deep-diving toothed whales, we found signatures of positive selection and molecular convergence affecting specific sites on *CDH23*, *TMC1,* and *SLC26A5*. Positively selected sites (PSS) were different in number, identity, and substitution rates (*dN*/*dS*) across riverine, coastal, and oceanic toothed whales.

**Conclusion:** Here we shed light on potential molecular mechanisms underlying the diversification of toothed whale echolocation. Our results suggest that toothed whale hearing genes changed under different selective pressures in coastal, riverine, and oceanic environments.

## BACKGROUND

Echolocation allows toothed whales to navigate and hunt underwater by producing high-frequency clicks and listening for the returning echoes [1]. It was a key development in toothed whale evolution, enabling their successful diversification across various aquatic environments, from shallow, murky rivers in South America to deep oceans in East Asia [1]. Although all toothed whales have echolocation, different species echolocate in very different environments, including rivers, and coastal and oceanic waters. These environments vary in physical conditions affecting sound propagation (e.g., depth, temperature, and salinity) [2], noise (i.e., other sounds produced by nearby sources), and clutter levels (i.e., echoes produced by a sonar when it interacts with objects other than the target), creating different selective pressures to the operation of dolphin sonars. For example, in shallow habitats such as rivers, estuaries, and coastal waters, sound waves encounter more obstacles and generate increased clutter, limiting sonar propagation to short distances [3]. Such constraints have likely favored the evolution of short-range, high-frequency sonars in riverine and coastal dolphins [3–9]. Oceanic dolphins, on the other hand, are found in deeper waters which are mostly noise-limited, meaning that overlapping sounds produced by other animals or geological phenomena are the primary obstacle to sonar propagation [3]. Accordingly, off-shore and pelagic-toothed whales usually employ sonars with lower frequencies and higher source levels that facilitate target detection over greater distances [2, 3]. The sonar differences between riverine, coastal, and oceanic species suggest that environmental conditions had an important role in shaping toothed whale echolocation.

Among mammals, high-frequency hearing is largely mediated by the inner and outer hair cells of the auditory cochlea (Figure 1). In mammalian outer hair cells (OHC), sound amplification and frequency tuning are mediated by the electromotile activity of prestin, encoded by the gene *SLC26A5* [10, 11]. The main reception of sound occurs in the inner hair cells (IHC), where the mechanosensory transduction (MT) complex converts mechanical stimuli from sound waves into electrochemical signals to the brain [12]. This complex includes a mechanosensitive ion channel that opens in response to tension from extracellular filaments (tip-links), producing the mechanotransduction current: an ion flow that depolarizes the hair cell towards releasing neurotransmitters, thus promoting sound perception [13, 14]. Two genes, *TMC1* and *CDH23*, encode major structural components of the MT channel and tip-links and are therefore essential to maintaining MT current. The transmembrane channel-like protein *TMC1* forms the pore region of the MT channel and is also involved in hair cell maturation and survival [15, 16]. *CDH23* encodes a calcium-dependent cell adhesion protein called cadherin-23, which forms the upper end of the tip-links [17] and is required for stereocilia movement, organization, and development [18, 19]. The depolarization of inner and outer hair cells is strongly dependent on their ionic gradient with the surrounding fluid, which is maintained by a series of adhesion proteins including claudin-14, encoded by the gene *CLDN14* [20]. Recent knockout experiments have shown that the absence of *CLDN14* causes IHC degeneration that affects the neurotransmission between hair cells and auditory nerves, suggesting an important role for this gene in the early processing of sound [21].

**Figure 1.**
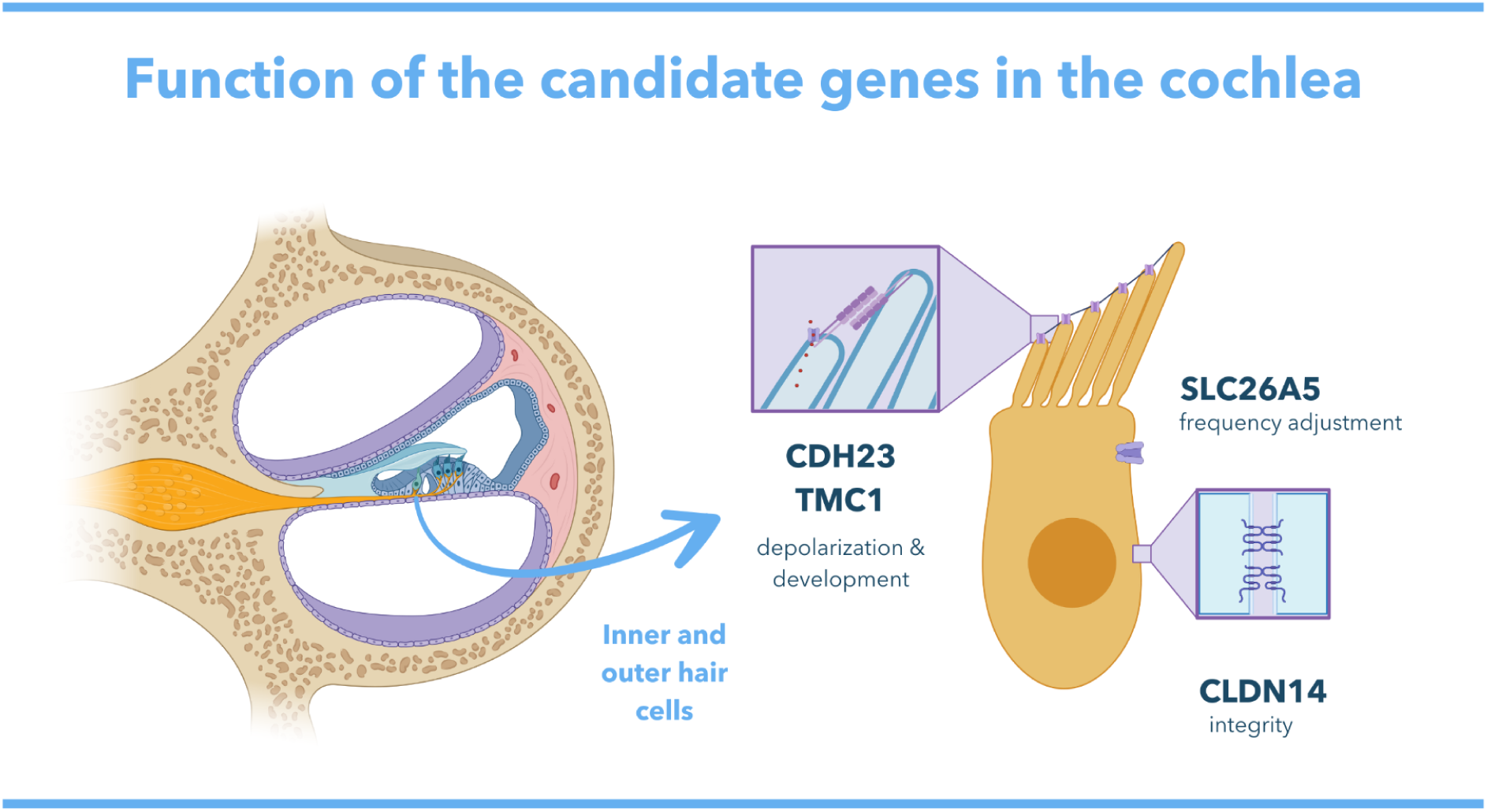
Graphic depiction of the cochlea on a longitudinal cut, showing the basilar membrane where the inner and outer hair cells are situated. Enhanced detail shows an Inner Hair Cell with the locations where the proteins encoded by *CDH23*, *TMC1*, *SLC26A5,* and *CLDN14* are expressed (made using BioRender and Canva Pro).

Growing evidence suggests that several hearing genes are functionally important to echolocation and were targeted by natural selection in toothed whale lineages [22–24]. Hearing genes have experienced multiple events of positive selection in toothed whales, and convergent evolution among echolocating toothed whales and bats, making them excellent candidates to investigate the adaptation of the toothed whale sonar to different environments. CDH23, CLDN14, TMC1, and SLC26A5 have been previously found under positive selection in both dolphins and bats and are strongly associated with high-frequency hearing [22–24][25][26–28][22–24]. However, it is still unknown how these genes evolved under the pressures of varying aquatic environments during the secondary radiations of toothed whales. The increased availability of whale genomes and coding sequence data allows for this question to be addressed comparatively across multiple species, which will contribute to unraveling the origins and diversification of cetacean acoustic behavior, and their broad evolutionary history.

Here we selected four candidate hearing genes (*CDH23*, *CLDN14*, *SLC26A5*, *TMC1*) with strong evidence of functional roles in high-frequency hearing to address how echolocation diversified among toothed whales in different habitats. Using multiple evolutionary models, we performed a comparative analysis of these genes across all odontocete families to investigate patterns of molecular evolution among riverine, coastal, and oceanic lineages. We hypothesized that the selective pressures of each environment have shaped the molecular evolution of hearing genes. We found distinct signatures of positive selection among riverine, coastal, and oceanic lineages, thus revealing potential molecular mechanisms to the evolution of toothed whale sonar in different habitats.

## RESULTS

We used a combination of evolutionary models to investigate the selective pressures acting on hearing genes, comparing toothed whales from distinct habitats.

The coding sequences for all four candidate genes were recovered for the same 37 odontocete species and seven outgroups (five baleen whales and two Artiodactyla), making our datasets identical in species representation across all genes. Overall, the nucleotide and protein-coding genes we estimated with IQtree recovered the most accepted relationships for all cetacean families, however, the topology for internal nodes, especially within the Delphinidae family, did not always reflect the species tree[29, 30]. Some species of river dolphins (notably the older lineages: *Platanista*, *Lipotes*, *Inia*, and *Pontoporia*), as well as deep diving oceanic dolphins (*Kogia breviceps*, *Kogia sima,* and *Physeter catodon*) showed particularly longer branch lengths, which could suggest evolutionary acceleration. *Platanista gangetica*, in particular, had very long branch lengths and was placed between Ziphiidae and Kogidae + Physeteridae on *CDH23*, as the outgroup for all toothed whales on *TMC1* and *SLC26A5*, and as ancestral to all cetaceans, on *CLDN14* (Figure S1-S7).

### Toothed whale lineages from different habitats show distinct evolutionary signatures

We found evidence of branch-wise positive selection in some coastal toothed whales for the genes *CDH23*, *SLC26A5*, and *TMC1*. Specifically, five out of the 13 coastal lineages have experienced episodic diversifying selection according to aBSREL, which was intense but restricted to only a few sites across the genes (Figure 2, Table S3). Overall, coastal lineages appear to have been under less constrained purifying selection compared with other cetaceans and artiodactyls, according to the ω estimates of the codeml branch models (Tables 2-4).

**Figure 2.**
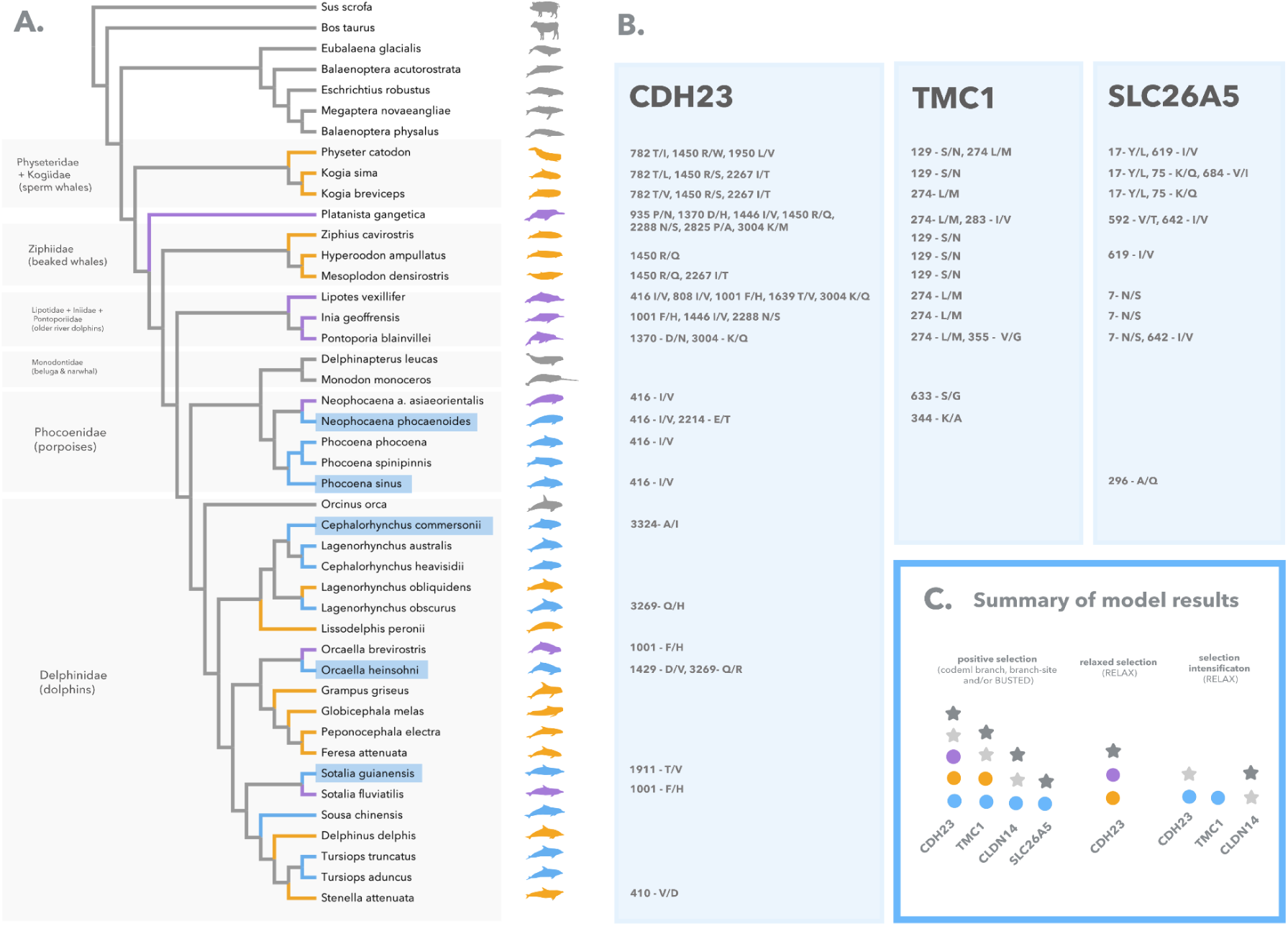
**A.** Phylogenetic tree of the toothed whale species included in this study, according to McGowen *et al.* (2020) [29]. Test branches are color-coded according to each environmental category/hypothesis: river = purple (H2a, H2b, H2c), oceanic = orange (H3), coastal/estuarine (H4) = blue; background branches are marked in gray. The five branches of coastal dolphins evolving under positive selection according to aBSREL are highlighted with blue rectangles. **B.** Robust Positively Selected Sites (PSS) for the genes *CDH23*, *TMC1*, and *SLC26A5*, using branch-site and site approaches, with the amino acid changes that were identified in the corresponding foreground branch. Rows in Figure 2B report the PSS that correspond to the lineages at the same level in Figure 2A. **C.** Summary of the selective regime found for each gene. Circles represent each environmental category (colored as in F2A), while stars denote evolutionary hypotheses: light grey for all extant toothed whales (H1b) and dark grey for ancestral toothed whales (H1a).

**Table 1.** Hypotheses were tested using the branch and branch-site codon models on codeml and HyPphy, along with the corresponding foreground branches, to investigate the evolutionary history of hearing genes associated with echolocation in toothed whales.

| Hypothesis | Foreground/test branches |
| --- | --- |
| 1a | Ancestral Odontoceti |
| 1b | Extant Odontoceti |
| 1c | All Odontoceti branches (extant + ancestors) |
| 2a | Riverine Odontoceti (*S. fluviatilis*, *N. a. asiaeorientalis*, *L. vexillifer*, *I. geoffrensis*, *P. gangetica*, *O. brevirostris*, *P. blainvillei*) |
| 2b | Phenotypically riverine Odontoceti (*L. vexillifer*, *I. geoffrensis*, *P. gangetica*, *P. blainvillei*) |
| 2c | Odontoceti that inhabit rivers (*S. fluviatilis*, *N. a. asiaeorientalis*, *L. vexillifer*, *I. geoffrensis*, *P. gangetica*, *O. brevirostris*) |
| 3 | Oceanic Odontoceti |
| 4 | Coastal Odontoceti |
| 5 | Riverine + Coastal vs. Oceanic Odontoceti |

**Table 2.**
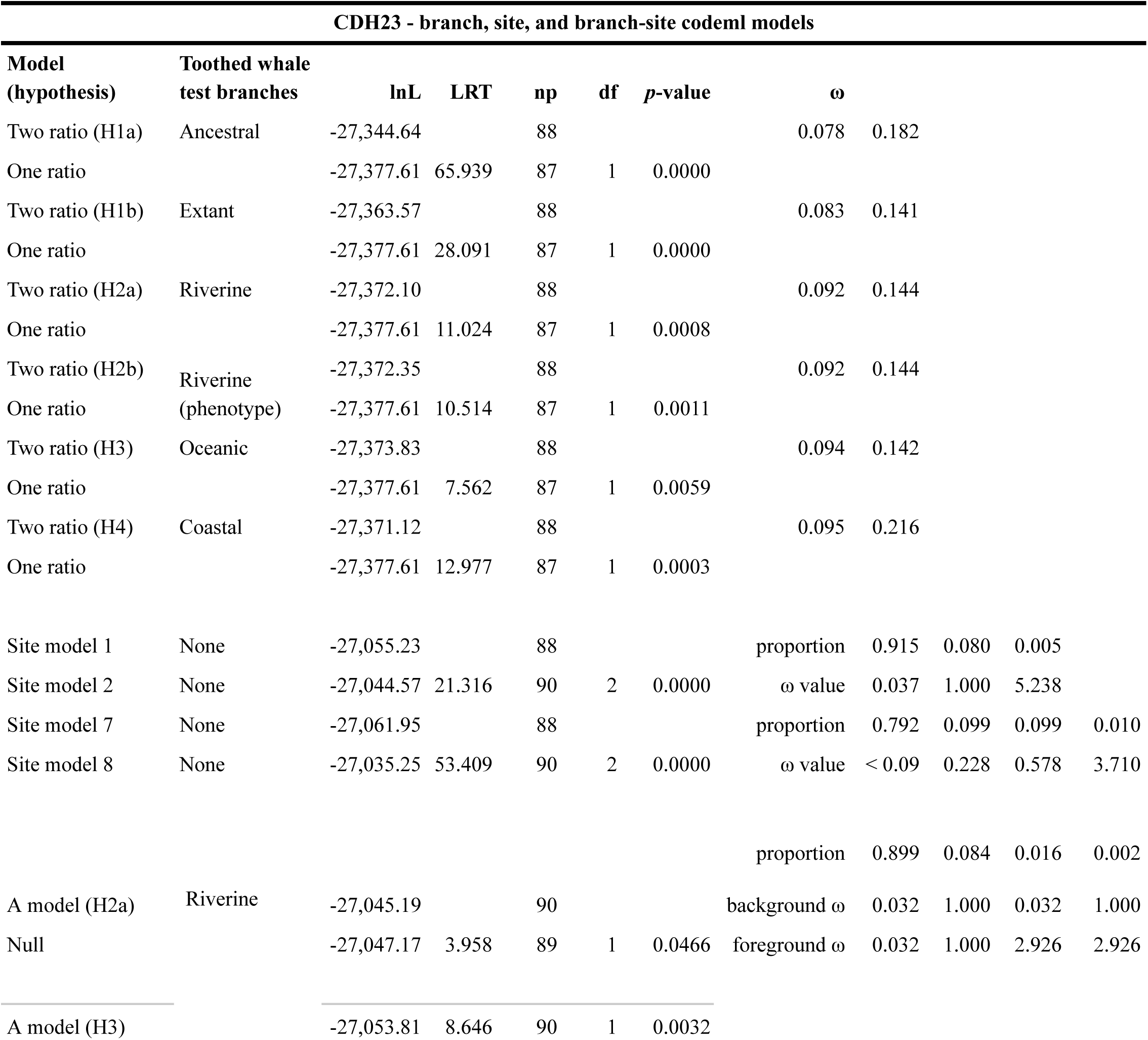
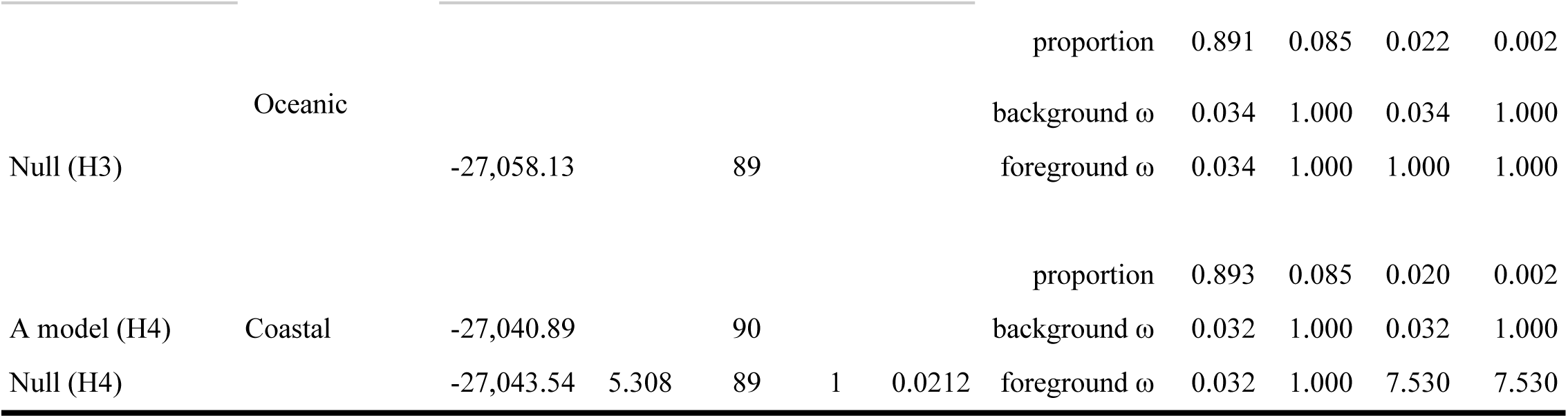
Codeml branch, site, and branch-site models with significant results for the gene *CDH23*. The hypotheses H1-H4 are outlined in Table 1. lnL = log likelihood; LRT = Likelihood Ratio Test; np = number of free parameters; df = degrees of freedom; ω = omega; A model = Alternative Model.

*CDH23* showed the strongest signatures of accelerated evolution, with three coastal lineages under episodic diversifying selection via aBSREL: *Cephalorhynchus commersonii*, *Sotalia guianensis,* and *Orcaella heinsohni* (Figure 2A, Table S3). Accordingly, the alternative branch and branch-site codeml models with coastal lineages as foreground branches had the best fit over the null models. This suggests that coastal toothed whales have experienced different selective pressures compared to other cetaceans, with a small proportion of their sites under intense positive selection (branch: LRT = 12.98, *p*-value = 0.0003; ω = 0.216; branch-site: LRT = 5.31, *p*-value = 0.0212, ω = 7.53) (Table 2). Finally, using RELAX, we also detected selection intensification on coastal branches when compared to extant toothed whales (Odontoceti), baleen whales (Mysticeti), and Artiodactyla (*Sus scrofa* and *Bos taurus*) (Figure 2C, Table S2). Among riverine and oceanic lineages, evidence of positive selection was also present for *CDH23* but was only supported by the two-model and the branch-site test for positive selection on codeml, suggesting that events of accelerated evolution among these lineages were weaker or fewer compared to coastal dolphins. More specifically, the codeml branch-site test estimated moderate positive selection in river dolphins (ω = 2.93), and neutral evolution in oceanic dolphins (ω = 1) (Figure 2C, Table 2). Furthermore, contrasting with coastal lineages, riverine and oceanic dolphins were under relaxed selection compared to other cetaceans and artiodactyls, according to RELAX (Figure 2C, Table S2).

*TMC1* had one lineage under episodic diversifying selection according to ABSREL: the Indo-Pacific finless porpoise (*Neophocaena phocaenoides*) (Figure 2A, Table S3). Similarly to *CDH23*, accelerated evolution among coastal toothed whales was supported by both branch and branch-site tests on codeml, with 0.36% of the sites under strong positive selection (branch: LRT = 4.92, *p*-value = 0.0264; ω = 0.515; branch-site: LRT = 14.31, *p*-value = 0.0001, ω = 149) (Figure 2C, Table 3). In addition, coastal dolphins experienced selection intensification relative to other cetaceans and artiodactyls, according to RELAX (Figure 2C, Table S2).

**Table 3.**
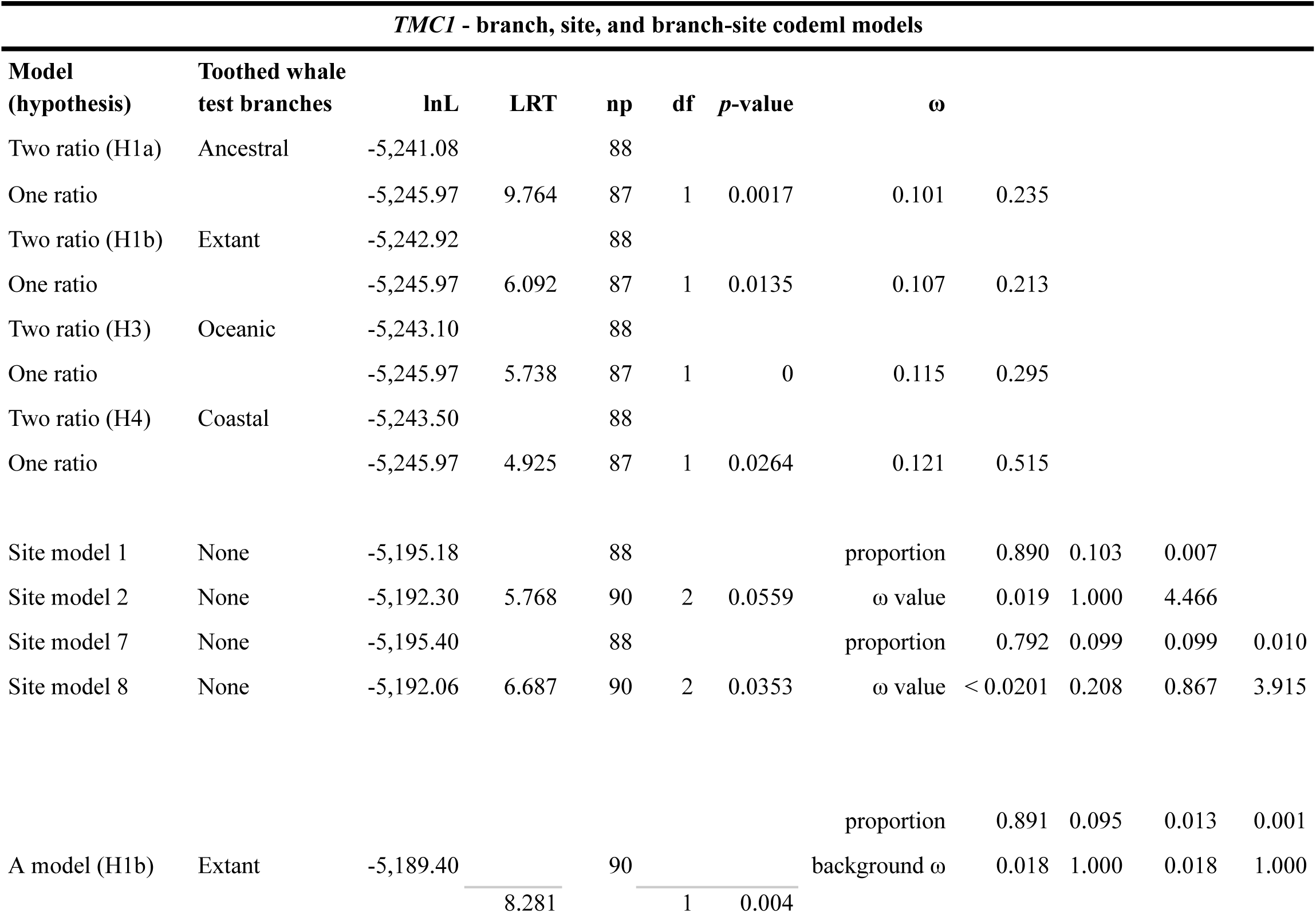
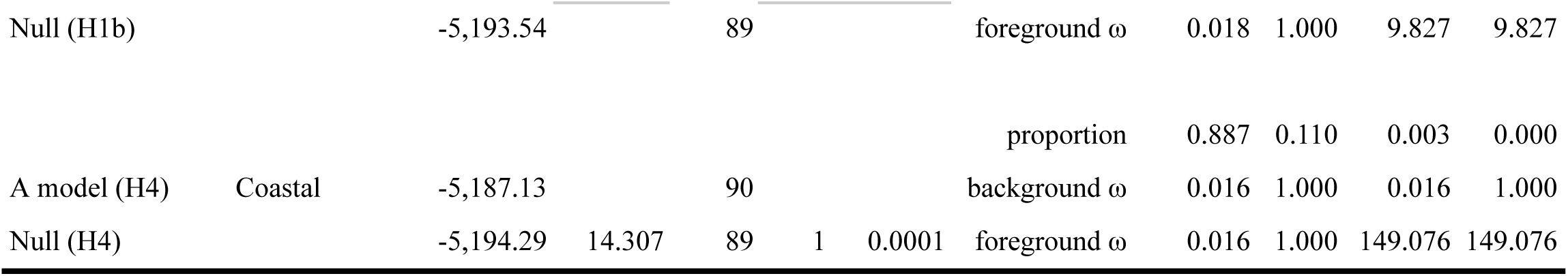
Codeml branch, site, and branch-site models with significant results for the gene *TMC1*. The hypotheses H1-H4 are outlined in Table 1. lnL = log-likelihood; LRT = Likelihood Ratio Test; np = number of free parameters; df = degrees of freedom; ω = omega; A model = Alternative Model.

For *SLC26A5*, we found one lineage enriched for episodic diversifying selection with aBSREL: the vaquita (*Phocoena sinus*) (Figure 2A, Table S3). Accelerated evolution among coastal toothed whales was identified with BUSTED and codeml branch-site test of positive selection, but not with the codeml branch model. The branch-site test estimated only 0.2% of the sites under strong positive selection (LRT = 4.60, *p*-value = 0.032, ω = 54.59) (Figure 2C, Table 4). Furthermore, no significant selection intensification was found with RELAX, suggesting that this gene is more conserved among toothed whales than the previous ones, with fewer episodes of positive selection (Figure 2C, Table S2).

**Table 4.** Codeml branch, site, and branch-site models with significant results for the gene *SLC26A5*. The hypotheses H1-H4 are outlined in Table 1. lnL = log-likelihood; LRT = Likelihood Ratio Test; np = number of free parameters; df = degrees of freedom; ω = omega; A model = Alternative Model.

| *SLC26A5* - branch, site, and branch-site codeml models |  |  |  |  |  |  |  |  |  |  |  |
| --- | --- | --- | --- | --- | --- | --- | --- | --- | --- | --- | --- |
| Model (hypothesis) | Toothed whale test branches | lnL | LRT | np | df | p-value | $\omega$ | | | | |
| Two ratio (H1a) | Ancestral | -5,770.21 |  | 88 |  |  |  |  |  |  |  |
| One ratio |  | -5,780.10 | 19.784 | 87 | 1 | 0 | 0.116 | 0.334 |  |  |  |
| Site model 7 | None | -5,726.33 |  | 88 |  |  | proportion | 0.747 | 0.093 | 0.093 | 0.066 |
| Site model 8 | None | -5,723.46 | 5.747 | 90 | 2 | 0.0564 | $\omega$ | < 0.09 | 0.143 | 0.272 | 1.737 |
|  |  |  |  |  |  |  | proportion | 0.869 | 0.129 | 0.002 | 0.000 |
| A model (H4) | Coastal | -5,722.90 | | 90 | | | background $\omega$ | 0.026 | 1.000 | 0.026 | 1.000 |
| Null (H4) | | -5,725.20 | 4.603 | 89 | 1 | 0.0319 | foreground $\omega$ | 0.026 | 1.000 | 54.593 | 54.593 |

Finally, for CLDN14, we found evidence for less stringent purifying selection affecting some coastal lineages according to the codeml branch test (LRT = 9.11, *p*-value = 0.0025, ω = 0.02).

We also investigated the selective pressures operating on the ancestral branches of toothed whales and among all extant toothed whales (Tables 2-4, Figure 2C). Our results suggest that diverse selective pressures have acted on extant and ancestral cetaceans across the four hearing genes. All genes showed some evidence of acceleration among ancestral branches or affecting all extant toothed whales, however, results were not consistent across multiple methods. Overall, extant toothed whales seem to be under more accelerated evolution compared to their ancestors and the outgroups (baleen whales + Artiodactyla), which is likely related to the positive selection affecting coastal species. For *CDH23* and *TMC1*, codeml branch models suggest that ancestral and extant cetaceans are under less constrained purifying selection compared to background branches (Table 2-4). On *CDH23*, we detected selection intensification on extant toothed whales compared to baleen whales + Artiodactyla, and relaxation of extant toothed whales relative to their ancestors. On *TMC1*, we found evidence of positive selection affecting extant toothed whales (Table 2-4). On *SLC26A5*, we found less constrained purifying selection on ancestral toothed whales, and positive selection on the branch leading to the sperm whales (*P. catodon*, *K.breviceps,* and *K.sima*) (Tables 4, S3). On *TMC1* and *CLDN14*, extant toothed whales had selection intensification compared to their ancestral branches (Table S2).

### Riverine, coastal, and oceanic dolphins have different sites under positive selection

To identify specific sites under positive selection in coastal, riverine, and oceanic toothed whales, we used a combination of sitewise approaches including the methods FUBAR, FEL, and MEME of the HyPhy package, the codeml site models M1, M2, M7 and M8, and the codeml branch-site model. Our analyses show that coastal, riverine, and oceanic toothed whales have different amounts and different sets of Positively Selected Sites (PSS) for the genes *CDH23*, *TMC1*, and *SLC26A5*, while *CLDN14* showed no sites under positive selection (Figure 2A, 3). Most PSS had changes that were exclusive to specific cetacean lineages from each environment, including the positively selected coastal dolphins identified with aBSREL (*C. commersonii*, *S. guianensis, O. heinsohni*, *N. phocaenoides*, and *P. sinus*), the older lineages of riverine dolphins (*I. geoffrensis*, *P. blainvillei*, *P. gangetica* and *L.vexillifer*) and deep-diving oceanic toothed whales (*P. catodon*, *K. sima, K. breviceps*). Overall, coastal and riverine toothed whales showed a higher number of PSS that were consistently found across multiple methods. Similar to the branch and branch-site analyses, we found the strongest signatures of accelerated evolution among coastal dolphins, especially for the gene *CDH23*.

**Figure 3.**
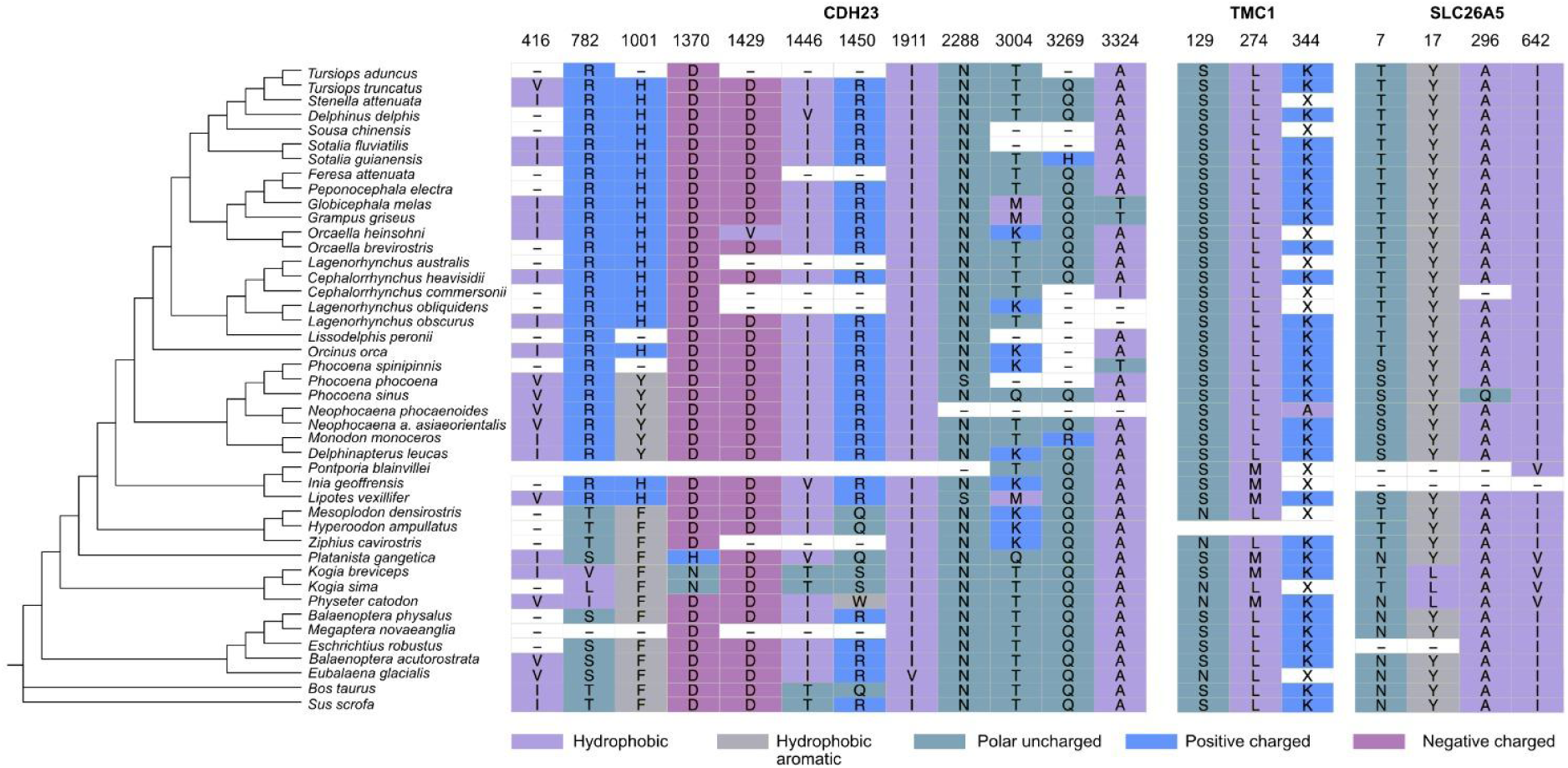
Aligned codon positions of Positively Selected Sites (PSS) showing amino acid changes on different odontocete species. Colors correspond to chemical differences between amino acids.

Coastal dolphins showed a total of eight PSS (CDH23: 6; TMC1: 1; SLC26A5: 1) with changes in seven species: *N.phocaenoides*, *P.phocoena*, *P.sinus*, *O.heinsohni*, *S.guianensis*, *L.obscurus*, *C.commersonii*. Most PSS had only one substitution affecting a single lineage at a time, except in two sites of CDH23: site 416, where the same substitution was found across all Phocoenidae (Figures 2-3, Tables S4-8), and site 2214, where both *N.phocaenoides* and *L.obscurus* have a substitution, however to different amino acids. Sites 1429, 1911, 3269, and 3324 of CDH23 and 344 of TMC1 showed strong signatures of positive selection, radical amino acid changes, and different evolutionary rates compared to riverine and oceanic dolphins (Tables S4-7). Three of these sites switched towards very hydrophobic amino acids (valine, isoleucine, and alanine), consistent with the radical changes in water solubility found with TREESAAP (hydropathy, solvent accessible reduction ratio, and surrounding hydrophobicity). Site 1429, in particular, changed from a hydrophilic to a completely hydrophobic amino acid (Asp - Val). Additionally, site 3269 also had a change affecting water solubility, however, in the opposite direction, from a neutral to a hydrophilic amino acid with a polar side chain (Gln - Arg), which is consistent with the radical change in isoelectric point found by TREESAAP. These similar changes in amino acid properties across independent lineages of coastal toothed whales (O.brevirostris, S.guianensis, C.commersonii), which were also enriched for branch-wise positive selection, could represent an emergent case of molecular convergence. Interestingly, most sites on CDH23 with radical property changes in riverine and oceanic dolphins also switch towards a decrease in hydrophily, either moving from hydrophilic to neutral, or from neutral to hydrophobic (1370: Asp - His, 1639: Thr - Val, 3004: Lys - Met). The only exception is site 410, which has an opposite change, from a very hydrophobic to a very hydrophilic amino acid (Figure 3).

River dolphins had the highest number of PSS across the three environments, with 11 sites identified in CDH23 alone, 4 sites in TMC1, and 3 sites in SLC26A5, with changes affecting all seven species (Tables S4-8). Some PSS had identical substitutions that happened in more than one independent lineage, indicating potential events of molecular convergence (CDH23: 416, 1001, 1446, 2288, and 3004; TMC1: 17, 642; SLC26A5: 274) (Figures 2-3).

Oceanic dolphins had nine PSS (CDH23: 9; TMC1: 3; SLC26A5: 4), with changes distributed across six species. There was a trend of identical substitutions among deep-diving species, which could indicate shared ancestral adaptations (when they happened in all members of a monophyletic group, such as site 782 of *CDH23* for *K. sima, K. breviceps* and *P. catodon*) or molecular convergence (when independent lineages evolved the same substitution, such as site 129 of *TMC1* for *P. catodon*, *K. sima*, *Z. cavirostris*, *H. ampullatus*, and *M. densirostris*) (Figures 2-3).

We also compared the selective pressures acting on specific sites across odontocete lineages. Contrast-FEL recovered sites with significantly different ω ratios in pairwise comparisons between riverine, coastal, and oceanic dolphins, indicating that some sites in the genes *CDH23*, *TMC1*, and *SLC26A5* have evolved under different selective pressures and natural selection rates in each environment (Table S11). Importantly, most of these sites were also under positive selection and showed radical amino acid changes on TREESAAP (Figure 3, Tables S4-8), suggesting that the substitution events in these sites might have resulted in important functional changes in the protein (Figure 4).

**Figure 4.**
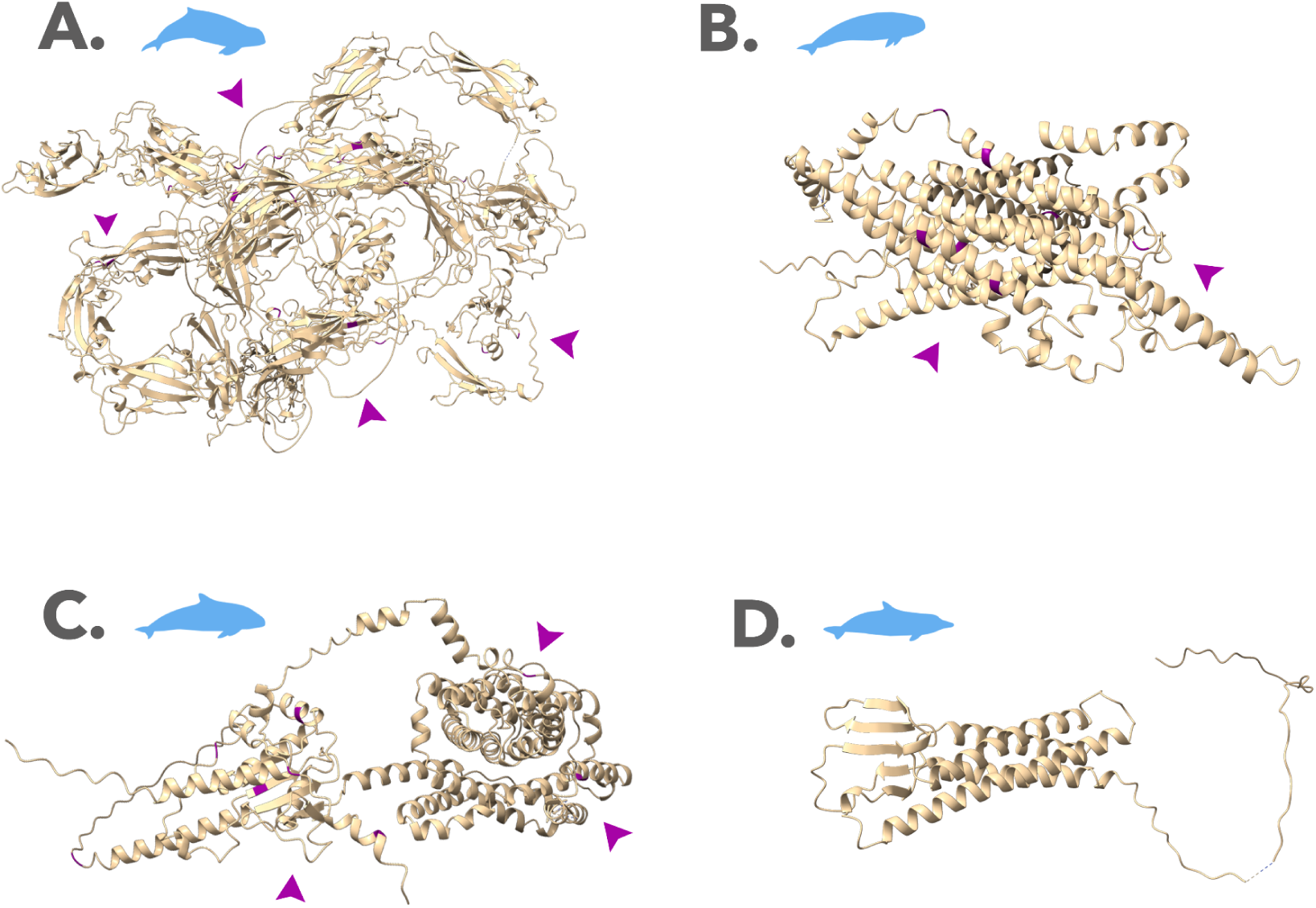
3D protein structures of the hearing genes: **A.** *CDH23* for *O.heinsohni*, **B.** *TMC1* for *N.phocaenoides*, **C.** *SLC26A5* for *P. sinus*, **D.** *CLDN14* for *S.guianensis*. PSS are highlighted in purple in the protein structure, and the approximate locations of each cluster of PSS are indicated with purple arrows.

All PSS were reported by at least two different sitewise methods and most (> 90%) were not repeated across environmental groups, suggesting that dolphin lineages from different habitats have undergone distinct events of sitewise positive selection (Figures 2-3). When all dolphins were selected as foreground branches, we found 18 PSS for *CDH23*, 6 for *TMC1*, and 6 for *SLC26A5*, and no PSS for *CLDN14*. Most of these sites corroborated previous signatures of positive selection found in each of the environmental categories, however, a few sites were only significant with all dolphins as foreground (Supplementary data).

## DISCUSSION

Echolocation and high-frequency hearing facilitated the adaptive radiation of toothed whales through a diversity of habitats with different selective pressures [3]. Here we show that the coding sequences of four hearing genes with a role in echolocation (*CDH23*, *SLC26A5*, *TMC1*, and *CLDN14*) have distinct signatures of molecular evolution among riverine, coastal, and oceanic species. This suggests that the different evolutionary constraints of each habitat have shaped the genetic diversity underlying the toothed whale sonar. Our comparative analysis across 37 odontocete species has revealed patterns of accelerated evolution within coastal and riverine lineages, supporting the hypothesis that these habitats generate specific selective pressures to sonar propagation and foraging, which are not found in the ocean [3, 5, 31–33]. We also found accelerated evolution in deep-diving oceanic toothed whales, such as sperm whales, which are known to have sonar and behavioral adaptations to hunt in great depths [34]. Furthermore, dolphins from different environments seem to have experienced positive selection in different sites, with a few cases of potential convergent evolution among riverine and deep-diving oceanic species. Finally, we found signatures of ancestral positive selection and accelerated evolution that could reflect the functional importance of these genes to cetaceans.

### Toothed whale lineages from different habitats show distinct evolutionary signatures

#### Coastal lineages

Our findings of positive selection and selection intensification in coastal lineages suggest that the coastal environment posed strong selective pressures on the evolution of hearing genes. The five lineages experiencing episodic diversifying selection (Figures 1, 2) are known to inhabit shallow waters (< 200 m deep) and are found in both coastal and estuarine environments, which interface with rivers (Figure 5, Table S1). Additionally, all species except for *P. sinus* are known to enter rivers, either occasionally (*S. guianensis*, *C. commersoni*, *O. brevirostris*) or frequently (*N. phocaenoides*)[35]. Shallow habitats with low visibility and high particle density are more acoustically complex than oceanic waters, and prey detection in these environments is usually limited by clutter (unwanted echoes from surrounding objects)[1, 3]. These restrictions create specific pressures that have likely been selected for short-range, high-frequency sonars in riverine and coastal dolphins [4, 5, 7, 9, 36]. This type of sonar is also advantageous for capturing small prey, which seems to be the preference of all positively selected lineages we report here [35].

**Figure 5.**
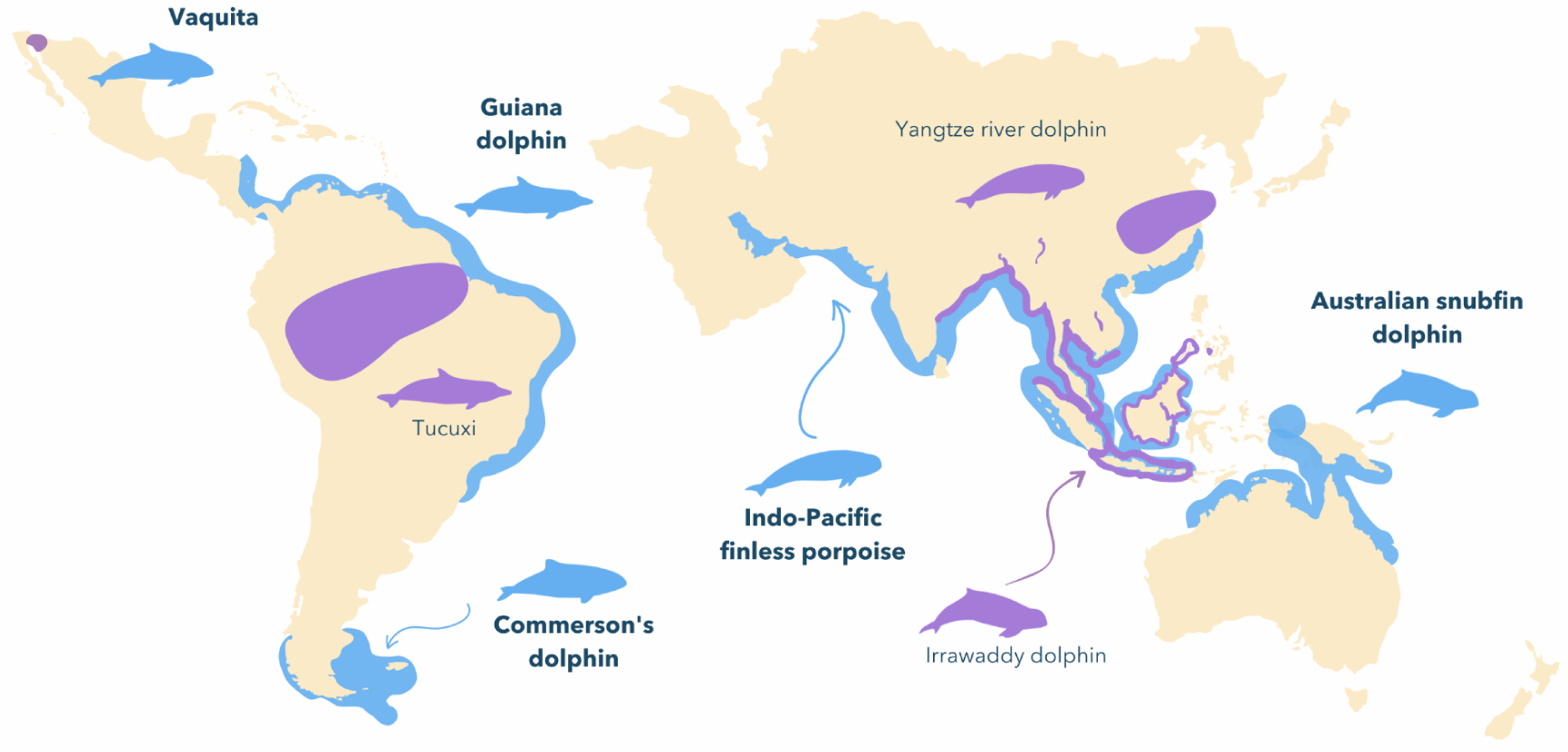
Distribution areas of the five lineages of coastal dolphins under positive selection (blue) and their riverine sister species (purple), as reviewed by Berta et al. (2015) [40].

Three of the five lineages under positive selection have recently diverged from their riverine sister species (*S. guianensis* and *S. fluviatilis*: 1.9 to 2 mya; *O. heinsohni* and *O. brevirostris*: 2.9 to 4 mya; *N. phocaenoides* and *N. asiaeorientalis asiaeorientalis*: 0.22 mya) [29, 37, 38] (Figure 5). Previous studies have detected differences in the sonar of these sister species, possibly related to habitat variation. For example, de Freitas et al. (2018) found higher source levels and centroid frequency in *O. heinsohni vs. O. brevirostris.* Another study revealed that the coastal Guyana dolphin (*S. guianensis*) had a larger bandwidth at the same dominant frequency when compared to its riverine sister species, the tucuxi (*S. fluviatilis*) [31]. Interestingly, the same study also recovered striking similarities between the tucuxi and the Amazon River Dolphin (*I. geoffrensis*), which could be correlated to their evolution in the Amazon waters [31]. Differences in the mean click parameters (e.g., frequency, bandwidth, interclick interval) between a coastal and a riverine subspecies of *N. phocanoides* (*N. a. asieorientalis and N. p. sunameri*) have also been detected, however, the authors caution that they could be due to a methodological limitation [39].

#### Riverine lineages

We uncovered signatures of convergence - i.e., identical substitutions in independent lineages - among the PSS of river dolphins, which is an important indicator of molecular adaptation (Figures 2B, 3) [41]. The oldest lineages of river dolphins (*Inia, Lipotes*, *Platanista*, and *Pontoporia*) share extensive morphological convergence associated with their independent colonization of freshwater habitats [42]. This convergence likely extends to acoustic phenotypes, since short-range, high-frequency, low-energy sonars have been reported in four species of river dolphins (*Neophocaena a. asiaeorientialis* - [43]; *P. gangetica* and *O. brevirostris* - [5]; *I. geoffrensis* - [4]. In this context, the signatures of accelerated evolution could represent a potential molecular basis for the convergent acoustic phenotypes of river dolphins. Interestingly, almost all PSS were found among the older river dolphins, consistent with their earliest colonization of the freshwater habitat [42].

#### Oceanic lineages

Finally, we found PSS exclusive to oceanic species, mainly sperm whales (*P. catodon*, *K. sima,* and *K. breviceps*) and beaked whales (*H. ampullatus* and *M. densirostris*), known for their deep diving behavior. These cetaceans have specific adaptations for hunting in deep waters and, most importantly, a long-range sonar to search for widely dispersed prey [3, 34, 44, 45]. In this study, we found potential signatures of convergent and ancestrally inherited substitutions among these deep-diving species, suggesting multiple evolutionary routes for their long-range sonars. The existence of a shared molecular basis to the deep diving sonar is corroborated by previous studies: positive selection was found in the ancestral lineages to Kogiidae + Physeteridae for CDH23 and SLC26A5, suggesting some adaptation for hearing in the deep ocean originated before the divergence of sperm whales [22]. On the other hand, there is also evidence for independent adaptation, as the cochlear morphology of sperm whales and beaked whales shows signatures of a rapid convergent evolution [46].

Surprisingly, no branch-wise positive selection was recovered for riverine or oceanic dolphins. However, the combined evidence of positively selected sites, with radical amino acid changes and altered evolutionary rates suggests that these dolphins were also subjected to environmental selective pressures. Additionally, the relaxed selection in *CDH23* affecting riverine, oceanic, and ancestral toothed whales might indicate that this gene experienced accelerated evolution caused by the relaxation of selective constraints, which could be associated with both adaptive and non-adaptive processes. Importantly, relaxed selection is a mechanism for evolutionary innovation and phenotypic plasticity, facilitating adaptation to new environments [47]. Therefore, the relaxed selection in *CDH23* could have allowed for an increased site-wise evolutionary rate, which led to the emergence of adaptive substitutions favored by positive selection.

Our results agree with previous findings that dolphins evolved specialized sonars and cochlear morphologies in each habitat [32, 33, 46]. Three types of environments, in particular, were associated with the strongest selective pressures: rivers, shallow coastal waters, and deep oceanic waters. The accelerated evolution of hearing genes among coastal and riverine dolphins could be related to their diversification in clutter-limited habitats [3–5]. In fact, the evolution of higher frequencies in riverine and coastal dolphins as a response to clutter-limitation could be related to the positive selection we found on CDH23, TMC1 and SLC26A5 since these genes are important to high-frequency hearing and frequency adjustment [27, 48, 49]. However, the striking differences in selection intensity between recently diverged coastal and riverine species within *Sotalia*, *Orcaella,* and *Neophocaena* suggest that coastal species experienced a more accelerated adaptation. Coastal dolphins, especially species inhabiting estuarine waters at the interface with rivers, are exposed to more dynamic environmental conditions, moving through a gradient of depth, salinity, and spatial complexity - which could result in more challenging selective pressures. Additionally, adaptation to environmental change could have been accelerated if selection acted upon ancestrally segregated variability, as occurred with the habitat-driven population structure of coastal bottlenose dolphins [50, 51]. Since the ancestors of *Sotalia*, *Orcaella,* and *Neophocaena* were likely coastal, they might have accumulated changes advantageous to this environment even before the separation of the sister riverine lineages. Future investigations focusing on the origin and phylogeography of recently diverged toothed whales that colonized different environments could reveal the exact conditions responsible for this accelerated evolution.

### Riverine, coastal, and oceanic dolphins have different sites under positive selection

We uncovered strong signatures of accelerated evolution in *CDH23*, *TMC1*, and *SLC26A5*, however, the sites targeted by positive selection in each gene largely varied among coastal, riverine, and oceanic toothed whales. Our results are consistent with previous studies that found an enrichment of positive selection events in cetacean hearing genes and molecular convergence in *CDH23*, *TMC1*, and *SLC26A5* among echolocating dolphins and bats [22, 27, 48].

*CDH23* showed the strongest signatures of adaptive evolution (i.e., lineages under positive selection, PSS, molecular convergence, and sites with different evolutionary rates among coastal, riverine, and oceanic dolphins). *CDH23* mediates the mechanotransduction in cochlear hair cells and has a variant involved in the Usher syndrome 1D, which causes congenital sensorineural deafness in humans [19]. *CDH23* has been previously found under positive selection among cetaceans in multiple ancestral lineages (Delphinidae, Delphinidae + Phocoenidae + Monodontidae + Lipotes + Inioidea, Kogiidae + Physeteridae), as well as under convergent evolution among marine mammals and echolocating mammals [22, 52–54]. Positive selection in *CDH23* has also been recovered for the hippopotamus lineage and their ancestral with cetaceans, and at the ancestral branch to all mammals, reflecting the overall importance of this gene to mammalian hearing [22, 55]. Additionally, recent evidence shows that *CDH23* is targeted by regulating micro-RNAs that could be associated with frequency variation between two subspecies of echolocating bats [56]. Most PSS of *CDH23* were located within the cadherin domains, near the Ca^2+^ binding sites essential for the tip-link adhesive function and structural integrity of this protein [57, 58] (Figure 4). More specifically, 4 PSS were located within 20-10 amino acids of the Ca^2+^ binding sites, and 11 PSS were within less than 10 amino acids. Mutations affecting positions near or at Ca^2+^ binding sites have been shown to disrupt the stability of cadherin domains and cause progressive hearing loss or even deafness [57] (Figure 4). Furthermore, 3 sites were located very close (within ≤ 10 amino acids) to conserved regions, more likely to be functionally important. The remaining PSS were in the cytoplasmic domain, responsible for anchoring the cadherin in the tip of the hair cell stereocilia (Figure 4). Finally, our TREESAAP analysis indicated radical changes in amino acid properties involving water solubility, suggesting that this was an important amino acid property targeted by natural selection. Therefore, decreases in water solubility affecting different sites could represent a potential molecular mechanism underlying the adaptation of dolphin hearing to different environments.

*TMC1* is a major component of the pore of mechanosensory channels in the inner ear, and therefore essential to the sensory transduction of sounds [15]. Mutations in this gene’s coding sequence have been associated with dominant and recessive non-syndromic hearing loss (DFNA36 and DFNB7/B11) [59, 60]. TMC1 also seems to have played a role in the early differentiation of the toothed whale sonar, with some evidence of positive selection on the ancestral branch of Phocoenidae + Monodontidae [22]. Previous evidence of positive selection in mammals and convergent evolution among echolocating dolphins and bats has linked *TMC1* to high-frequency hearing for echolocation [48, 55]. Site 129, with a substitution from serine to asparagine, was found here under positive selection in deep-diving sperm whales and beaked whales and was previously reported under adaptive convergence between two species of echolocating bats, *Rhinolophus ferrumequinum* and *Pteronotus parnellii* [48]. Positively selected sites in *TMC1* are mapped onto different protein domains, mostly cytoplasmic, except for sites 274 and 283, which are immediately preceding or within the transmembrane helix domain 2 [15] (Figure 4). On the 3D structure of *TMC1*, mapped for *C.elegans*, these sites are lining up the pore walls and interacting with chemical ligands, such as cholesterol and undecan-1-ol [61]. On *C.elegans*, site 344 interacts directly with CALM, a calcium-binding homologous of the vertebrate CIB2, which modulates the activity of the mechanosensory transduction complex and mediates the interactions between *TMC1* and other proteins of the complex [61]. Site 633 is of particular interest since it appears along the pore walls and interacts with the CALM protein, and chemical ligands [61]. Therefore, although not situated directly in the pore-forming transmembrane domains [15], some of the positively selected sites found here are potentially important to the functional activity of *TMC1*, given their location at the surface of the pore walls and interaction with other proteins in the mechanosensory complex.

*SLC25A5* or prestin is the mediator of OHC electromotility, which promotes frequency adjustment during cochlear amplification. It is also important for OHC survival and maintenance of local afferent and efferent circuits [10, 62]. Mutations in prestin underlie autosomal recessive deafness (DFNB61) [63], and extensive evidence for positive selection and convergence between dolphins and bats implicate a fundamental role for this gene in echolocation, especially regarding frequency adjustment [24, 27, 28]. Prestin has also been found to be under selection in several other mammalian branches, including the ancestors of toothed whales, mammals, and tetrapods, which is consistent with this gene’s essential role in hearing [22, 54, 64]. Here we found several events of positive selection within the coding sequence of prestin, all within cytoplasmic domains, more specifically the N terminal domain and the STAS domain. Although these are not the main active domains responsible for the cell’s motor activity, they have been shown as critical for protein dimerization [65].

Furthermore, many of the PSS we report here have also been previously found in studies investigating the molecular convergence between echolocating toothed whales and bats [24, 26]. For example, here we found the PSS 17, 75, and 642 showing the same substitutions as reported by Li et al. (2010), but in different cetacean branches. Additionally, the PSS 7 and 592 found here were also reported by Li et al. (2010), but with different substitutions in different branches. Interestingly, for site 619, we recovered the same substitution from Isoleucine to Valine occurring in the same deep-diving toothed whale branches (*P. macrocecphalus* and *H. ampullatus*) as reported by Liu et al. (2010). Additionally, almost all positively selected branches found by both our study and Liu et al. (2010) are oceanic deep-diving toothed whales that employ extreme frequencies in their echolocation: sperm whales and beaked whales with very low frequencies, and dwarf/pygmy sperm whales with very high frequencies. Given the role of prestin in frequency adjustment, substitutions that increase protein stability could be adaptive for cetaceans as a whole, but especially for species that vocalize in extreme frequencies. This is consistent with the fact that the ancestor of baleen whales, which are extremely low-frequency vocalizers, also shows the I-V substitution on site 619. Site 619 could therefore have evolved convergently in echolocating bats and dolphins, as well as baleen whales, as an adaptation to hearing extremely high or low frequencies.

*CLDN14* showed no signatures of positive selection or convergence at the site level. However, we did find branch-wise positive selection and selection intensification in ancestral toothed whales, similar to Xu et al. (2013) [66], and across all toothed whales. This suggests that *CLDN14* was important to the early development of echolocation, and although probably not related to environmental adaptation, was under important selective pressures among toothed whales.

The collective findings of our study, together with previous evidence of ancestral positive selection [22] and dolphin-bat convergence on CDH23, SLC26A5 and TMC1 [27, 48, 54, 67], suggest that these hearing genes have been involved in multiple events of adaptation in toothed whales, as well as other echolocating (and non-echolocating) mammals. This is expected, given the overall importance of these genes to mammalian hearing and frequency modulation. However, the different patterns of positive selection and substitutions affecting hearing genes in each environment suggest they are subject to diverse selective pressures across space and time. Together with McGowen *et al.* (2020)’s findings, our results suggest that CDH23, SLC26A5, and TMC1 were important to toothed whale evolution both at ancestral stages, for the development of the sonar, and during the speciation of extant lineages and adaptation to different habitats in the secondary radiations.

### Limitations

Here we investigated the molecular evolution of hearing genes using a comprehensive dataset of toothed whale species to investigate how their sonars could have changed in riverine, coastal, and oceanic environments. To our knowledge, this is the first study to uncover a molecular basis for the divergence of toothed whale echolocation in different habitats. Previous studies that found positive selection in cetacean hearing genes focused on their transition from terrestrial to aquatic habitats and did not include a comprehensive set of dolphin species. Most of these studies have shown robust evidence for convergence between dolphins and bats when including only a few species of cetaceans, however, we observed that the pattern of identical substitutions they found does not extend to all cetacean species. This further supports the hypothesis that echolocation might have diversified in toothed whales following multiple adaptive pathways, which could have been influenced by a collection of environmental factors [3]. The evolution of a complex phenotype such as echolocation is likely to involve a combination of selective pressures, which can result in repeated substitutions under a convergent environment or lineage-specific changes that could reflect their particular evolutionary history. In this study, we found molecular evidence for both patterns, suggesting that the diversification of echolocation across extant toothed whales was a more complex process than originally thought.

The patterns of accelerated evolution we report here might also be correlated to other evolutionary scenarios unrelated to environmental differences. The toothed whale sonar is a complex feature influenced by multiple different factors, including allometric scaling, prey size, and ecological interactions. For example, three of the coastal lineages under diversifying selection (*N. phocaenoides*, *P. sinus*, *C. commersonii*) are Narrow Band High Frequency (NBHF) echolocators, meaning that they operate sonars within a 125-140 kHz peak frequency range with a narrow bandwidth (11-20 kHz) (Figure 3C) [68–70]. NBHF sonars are thought to have evolved independently in four extant odontocete families (Pontoporiidae, Kogiidae, Phocoenidae, and Delphinidae) under the selective pressures of predation by killer whales and other extinct raptorial toothed whales [70]. Therefore, our results could reflect a case of molecular convergence among NBHF species, which has not been yet reported. Furthermore, a part of current variation in the echolocation systems of toothed whales is likely due to phenotypic plasticity, and not underlying genetics. Dolphins can perform subtle adjustments in their biosonar parameters, such as frequency and output levels, in face of environmental changes such as moving to a deeper or more clustered area while foraging [71].

Our findings could also have been influenced by the pleiotropic effects of the studied genes, since they are potentially involved in other biological processes, such as vision and locomotion. However, current evidence from functional essays, knock-out experiments, and evolutionary studies shows a much larger and better-supported role of these genes in auditory processes.

## CONCLUSIONS

We investigated the evolution of the hearing genes *CDH23*, *TMC1*, *SLC26A5*, and *CLDN14* in toothed whale lineages from different habitats. We uncovered strong positive selection and accelerated evolution among coastal lineages, in addition to multiple sitewise changes with convergent and divergent patterns reflecting habitat distinctions. Our results suggest multiple pathways for the evolution of toothed whale echolocation and reveal a potential molecular basis for the sonar differences between riverine, coastal, and oceanic dolphins. Future studies focusing on the regulation and expression of these genes could bring further insights into the molecular changes underlying the diversification of the toothed whale sonar, and reveal additional genes functionally relevant to echolocation. The identification of genomic regions important to other echolocation processes, such as sound production and cognitive processing, will allow us to explore different hypotheses on how the toothed whale sonar changed under different selective constraints. Genome-wide scans indicate that there is a large pool of unexplored genes potentially correlated with cetacean adaptations [52, 72]. Comparing the evolution of these genes in a phylogenetic framework will advance our understanding of recent cetacean diversification, and the molecular processes underlying adaptation in different habitats.

## METHODS

### Sequence download and alignment

We selected four candidate hearing genes - *CDH23*, *SLC26A5*, *TMC1*, and *CLDN14* to investigate the molecular evolution of the toothed whale sonar. Previous studies on the molecular evolution of echolocating mammals suggest that these genes have a strong functional association with echolocation and high-frequency hearing, and not just to hearing in general. In addition, unlike other hearing genes, our candidate genes have reduced levels of pleiotropy and have been well-sequenced and characterized in toothed whale species. This makes them ideal candidates to investigate molecular changes specifically affecting toothed whale branches during the secondary radiations of cetaceans.

We retrieved the coding sequences for each of the candidate genes from the Nucleotide, Sequence Read Archive (SRA), and Genome databases in Genbank. By combining these three data sources, we included 42 species of cetaceans (37 toothed whales and five baleen whales - Table S12), along with two external Artiodactyla lineages: *Sus scrofa* and *Bos taurus*. SRA sequences were extracted from a concatenated alignment of 3191 coding genes [29], which was aligned against the sequences of *Orcinus orca* and *Tursiops truncatus* using MAFFT web-server 7, to recover cetacean orthologs for the four candidate genes [73]. Since the SRA data had higher levels of missing data compared to the other two, we kept only the species where more than 50% of the entire coding sequence of the gene was available. When a sequence was available on all three data sources, we selected the one with the most complete coding sequence. The genomic sequences of *Platanista gangetica* (GCA_017311385.1) and *Phocoena sinus* (GCF_008692025.1) were downloaded from the Genome database and coding sequences for the candidate genes were extracted using BLAST 2.12.0 [74], since Nucleotide or SRA sequences were not available for these species.

Additionally, we assembled the coding sequences of *Sotalia guianensis* and *Sotalia fluviatilis* from newly sequenced whole genomes, using BLAST to locate the sequences, a custom python script for extraction, and Aliview 1.27 [75] for alignment, assembly, and visual inspection of the gene fragments. Only the best hits of sequences (identities over 94%) were kept for alignment (Tables S9-S10).

### Phenotypic and environmental categories

To compare the evolutionary patterns of hearing genes among toothed whales from different habitats, we first assigned each species to an environmental category among “riverine”, “coastal”, and “oceanic”, according to their habitat preferences [40, 76, 77] (Figure 2). To avoid confounding effects in the association between molecular changes and different environments, when the species showed no predominant habitat, they were assigned to a fourth category termed “generalist”, which was not included in our tests. To assess the diversity of acoustic properties in toothed whale sonars, we used the classification proposed by Jensen et al. (2018) [3] and assigned each odontocete species to one of the four sonar categories described in the study (Table S1).

### Sequence alignment and phylogenetic reconstruction

We assembled the coding sequences into multi-species alignments with the MAFFT web server, applying the G-INS-i alignment strategy (with the remaining settings on default) [73]. We translated the alignments into amino acids using Aliview [75] and used PAL2NAL [78] to obtain the codon alignments used in all selection analyses. We used W-IQ-TREE multi-core 1.6.12 [79] to reconstruct maximum likelihood gene trees for each multi-species codon alignment under the GY+F+G4 model [80]. To evaluate branch support of the consensus trees, Ultrafast Bootstrap Analysis was performed with 10,000 bootstrap replicates, 1,000 maximum iterations, and a minimum correlation coefficient of 0.99[81]. The single branch tests SH-aLRT (with 5000 replicates) and Approximate Bayes were conducted to further assess support and maximize tree confidence [82, 83].

### Selection analyses

To evaluate the role of selective pressures in the evolution of the four hearing genes, we investigated signals of positive selection, substitution rate shifts, and selective constraints on branches and sites using a combination of codon models implemented on HyPhy 2.5 [84] and PAML 4.9 [85]. These codon models are based on the estimation of the ω (*dN* / *dS*) value, the ratio of nonsynonymous to synonymous substitutions, which is used to infer the strength of selection on codon alignments along single branches (branch-models), sites (site-models), or both (branch-site models). An ω > 1 indicates a higher accumulation of non-synonymous substitutions, which is interpreted as Darwinian positive selection. An ω < 1 indicates a higher rate of synonymous substitutions and it is interpreted as negative or purifying selection. Finally, an ω value equal to or close to 1 indicates similar rates of non-synonymous and synonymous substitutions, consistent with neutral evolution.

For each method, we explored different sets of hypotheses to compare the evolution of the four hearing genes among different cetacean lineages. The foreground branches for each hypothesis are described in Table 1. These hypotheses were designed to compare the evolutionary rates between (1) the Odontoceti ancestor *vs.* extant lineages, (2) all extant Odontoceti *vs.* lineages from a specific habitat - i.e., riverine, coastal, or oceanic, and (3) lineages from different habitats with each other (Table 1). Additionally, we specified hypotheses 2a, 2b and 2c to compare the oldest lineages of river dolphins (*L.vexillifer*, *I.geoffrensis, P. gangetica* and *P.blainvillei*), which share convergent morphological features, with more recent lineages (*S. fluviatilis, N. a. asiaeorientalis, O. brevirostris*) that are also exclusive to freshwater habitats but do not show the “classic” river dolphin convergent morphology.

#### Branch-models

To obtain an initial assessment of positive selection signatures among the four hearing genes, we used BUSTED (Branch-Site Unrestricted Statistical Test for Episodic Diversification) [86] with different branch partitions corresponding to our evolutionary hypotheses (Table 1). Then, we used the codeml one-ratio model and two-ratio model [87, 88] to test for different evolutionary patterns among coastal, riverine, and oceanic dolphins. The one-ratio model is a commonly used null hypothesis for codon evolution, which specifies a single ω value for all branches. The two-ratio model, on the other hand, estimates different ω values to foreground and background branches and also allows for multiple categories of foreground branches, each one with a distinct ω. We specified the foreground branches on this test to explore hypotheses 1a, 1b, 2a, 2b, 3, 4, and 5 (Table 1). For each hypothesis, a likelihood-ratio test (LRT) was performed to compare the fit of model 0 versus model 2 with 2ω [87, 88]. Next, we used RELAX [89] as a complementary test to explore the variation in evolutionary rates across odontocete species from riverine, coastal, and oceanic habitats, with foreground branches specified according to hypotheses 1a, 1b, 2a, 3, and 4 (Table 1).

#### Branch-site models

To identify individual branches experiencing episodic diversifying selection on each gene, we used aBSREL (adaptive Branch-Site Random Effects Likelihood) [90] and the codeml branch-site test for positive selection [91]. Branch-site models allow for ω variations among both branches and sites and are therefore useful to investigate positive selection affecting specific sites on the foreground branches. Both methods estimate site rate classes with different ω values and perform an LRT test between two models: an alternative model, where foreground branches have sites with ω > 1, and a null model, where ω < 1 is not allowed. However, aBSREL infers the optimal number of site rate classes for each gene and does not test for selection at specific sites.

We used aBSREL as an initial screening to identify specific lineages of riverine, coastal, or oceanic dolphins under positive selection, and then the codeml branch-site test of positive selection to search for specific sites under positive selection in those lineages. Positively Selected Sites were identified using the Bayes Empirical Bayes (BEB) and the Naive Empirical Bayes (NEB) methods on codeml, with a cutoff criteria at posterior probabilities > 0.9 [92]. aBSREL adjusts all *p*-values obtained from individual tests for multiple comparisons using the Bonferroni-Holm procedure, which controls the family-wise false positive rate [90, 93].

#### Site models

To further investigate the evolutionary history of each gene and compare the lineages from distinct habitats, we looked for signatures of positive selection in individual sites using the codeml site models M1a (neutral), M2a (selection), M7 (beta), and M8 (beta & ω), as well as the HyPhy methods FUBAR (Fast, Unconstrained Bayesian Approximation), MEME (Mixed Effects Model of Evolution), FEL (Fixed Effects Likelihood) and Contrast-FEL [94–97]. We considered as robust the Positively Selected Site (PSS) that was reported by at least two different methods, with the following cut-off values for statistical significance: posterior probability > 0.9 (codeml and FUBAR) and *p*-value < 0.1 (MEME, FEL, and Contrast-FEL).

### Amino-acid changes and protein structure modeling

We used TreeSAAP v.3.2 [98] to identify radical changes in amino acid properties associated with non-synonymous substitutions in each site, under a phylogenetic framework. TreeSAAP implements ancestral node reconstruction using the PAML program baseml [85] and compares sequences to infer amino-acid substitution events. These substitutions are classified into eight groups based on the magnitude of their physical-chemical effects, ranging from mild to radical substitutions. We used a goodness-of-fit test, which yields a z-score, to determine whether these mutations were potentially affecting the physicochemical properties of the amino acids. Results were treated using the software IMPACT-S v. 1.0.0 [99], where only sites with z-scores above the threshold and substitutions in the categories 6-8 were considered as under significant radical changes.

To further explore how sitewise changes could have potentially affected protein structure and function, we used AlphaFold2 as implemented in the ColabFold v1.5.2-patch [100] to build three-dimensional models of the *CDH23*, *TMC1* and *SLC26A5* and *CLDN14* proteins for the lineages under positive selection (*C. commersoni*, *N. phocaenoides, O. brevirostris*, *P. sinus* and *S. guianensis*). We then located the PSS in the 3D protein structure and retrieved functional and structural annotations from Uniprot to characterize the domains where PSS are located [101].

## LIST OF ABBREVIATIONS

IHC = Inner Hair Cell

NBHF = Narrow Band High Frequency

PSS = Positively Selected Sites

OHC = Outer Hair Cell

## DECLARATIONS

### Ethics approval and consent to participate

Not applicable.

### Consent for publication

Not applicable.

### Availability of data and materials

All sequence and alignment data and codes used for this project, as well as results files from each analysis, are available at: https://github.com/leticiamagpali/Magpali_2024_echolocation/

## Authors’ contributions

Author’s contributions have been described following the Contributor Role Taxonomy (CRediT).

LM: Conceptualization, Data curation, Formal analysis, Funding acquisition, Investigation, Methodology, Project administration, Software, Visualization, Writing – original draft, Writing – review & editing.

MFN: Conceptualization, Funding acquisition, Methodology, Project administration, Resources, Supervision, Visualization, Visualization, Writing – original draft, Writing – review & editing.

ER: Software, Supervision, Visualization, Writing – original draft, Writing – review & editing.

LF: Software, Writing – review & editing.

AP: Supervision, Software, Writing – review & editing.

## Acknowledgements

The authors would like to thank Ahmad Hamdan, Daniel Magpali for proof-reading the manuscript and Joseph Bielawski for excellent methodological suggestions.

