## Supplemental material for "Molecular evolution of toothed whale genes reveals adaptations to echolocating in different environments"

1. Laboratório de Genômica Evolutiva. Departamento de Genética, Evolução, Microbiologia e Imunologia, Universidade Estadual de Campinas (Unicamp), Campinas, São Paulo, Brasil
2. Department of Biology, Dalhousie University, Halifax, Nova Scotia, Canada

**Table S1.** Echolocation parameters for all species of toothed whales used in this study. Fp = peak frequency; BW-3db = Bandwidth; SLpp = peak to peak source levels; Duration = click duration in seconds; Type = echolocation type as classified by Jensen et al. (2018) [1]

| Species | **Habitat (predominant)** | **Echolocation parameters** | | | | |
|---|---|---|---|---|---|---|
| | | Fp | BW-3dB | SLpp | Duration | Type |
| *Stenella attenuata* | Oceanic | 69 | 83 | 212 | 43 | I - Delphinidae |
| *Tursiops aduncus* | Coastal | 124 | 62 | 205 | 16 | I - Delphinidae |
| *Tursiops truncatus* | Coastal | - | - | 200 | 22 | I - Delphinidae |
| *Delphinus delphis* | Oceanic | - | - | - | - | I - Delphinidae |
| *Sousa chinensis* | Coastal | 109 | 50 | 188 | 19 | I - Delphinidae |
| *Sotalia guianensis* | Estuarine and coastal | - | - | - | - | I - Delphinidae |
| *Sotalia fluviatilis* | Riverine | 93.1 | 21.5 | 201.8 | 93.1 | I - Delphinidae |
| *Feresa attenuata* | Oceanic | 70 | - | 210 | 30 | I - Delphinidae |
| *Peponocephala electra* | Oceanic | - | - | - | - | I - Delphinidae |

| | | | | | | |
|---|---|---|---|---|---|---|
| *Globicephala melas* | Oceanic | 50 | 46 | 196 | 23 | I - Delphinidae |
| *Grampus griseus* | Oceanic | 49 | 27 | 220 | 40 | I - Delphinidae |
| *Orcaella brevirostris* | Estuarine, riverine | 101 | 64 | 195 | 13 | I - Delphinidae |
| *Orcaella heinsohni* | Coastal and estuarine | 104 | 69 | 200 | 12 | ? |
| *Lissodelphis peronii* | Oceanic | - | - | - | - | I - Delphinidae |
| *Lagenorhynchus obscurus* | Coastal | 74 | 67 | 191 | 16 | ? |
| *Lagenorhynchus obliquidens* | Oceanic | - | - | - | - | I - Delphinidae |
| *Cephalorhynchus heavisidii* | Coastal | 125 | 15 | 173 | 74 | ? |
| *Lagenorhynchus australis* | Coastal | 126 | 15 | 185 | 92 | 2 - NBHF |
| *Cephalorhynchus commersonii* | Coastal | 132 | 21 | 177 | 78 | 2 - NBHF |
| *Orcinus orca* | Estuarine, coastal and oceanic | 29 | 25 | 203 | 41 | I - Delphinidae |
| *Phocoena sinus* | Estuarine and coastal | - | - | - | - | 2 - NBHF |
| *Phocoena spinipinnis* | Coastal | - | - | - | - | 2 - NBHF |
| *Phocoena phocoena* | Coastal | 137 | 15 | 186 | 62 | 2 - NBHF |
| *Neophocaena phocaenoides* | Coastal | 129 | 21 | 178 | 44 | 2 - NBHF |
| *Neophocaena a. asiaeorientalis* | Riverine | 129 | 22 | 176 | 48 | 2 - NBHF |
| *Monodon monoceros* | Arctic (coastal and oceanic) | 69 | 30 | 210 | 28 | I - Delphinidae |
| *Delphinapterus leucas* | Panarctic, coastal | 41 | 13 | 218 | - | I - Delphinidae |
| *Pontoporia blainvillei* | Coastal, estuarine | 139 | - | - | 212 | 2 - NBHF |
| *Inia geoffrensis* | Riverine | 96 | 50 | 190 | 14 | I - Delphinidae |
| *Lipotes vexillifer* | Riverine | - | - | - | - | I - Delphinidae |

| | | | | | | |
|---|---|---|---|---|---|---|
| *Mesoplodon densirostris* | Oceanic | 32 | 12.4 | 211 | - | III - Ziphiidae |
| *Hyperoodon ampullatus* | Oceanic | 54 | - | 203 | 276 | III - Ziphiidae |
| *Ziphius cavirostris* | Oceanic | 40 | 12 | 214 | 200 | III - Ziphiidae |
| *Platanista gangetica* | Riverine | 59 | 44 | 183 | 22 | I - Delphinidae |
| *Kogia breviceps* | Oceanic | 130 | 8 | - | 119 | 2 - NBHF |
| *Kogia sima* | Oceanic | - | - | - | - | 2 - NBHF |
| *Physeter catodon* | Oceanic | 12 | - | 240 | 120 | IV - Sperm whale |

**Table S2.** RELAX results for the genes with evidence of shifts (intensification or relaxation) in the strength of natural selection. K = selection intensity parameter. LR = likelihood ratio test.

| Gene | Foreground branches | Reference branches | Selection shift | K | p | LR |
|---|---|---|---|---|---|---|
| CDH23 | Extant odontocetes | Extant Mysticeti + Artiodactyla | intensification | 3.91 | 0.000 | 16.46 |
| | Extant odontocetes | Ancestral odontocetes | relaxation | 0.84 | 0.046 | 3.99 |
| | Riverine odontocetes | Extant Odontocetes + Mysticeti + Artiodactyla | relaxation | 0.69 | 0.000 | 22.04 |
| | Oceanic odontocetes | Extant Odontocetes + Mysticeti + Artiodactyla | relaxation | 0.67 | 0.003 | 9.12 |
| | Coastal odontocetes | Extant Odontocetes + Mysticeti + Artiodactyla | intensification | 1.57 | 0.000 | 14.32 |
| TMC1 | Coastal odontocetes | Extant Odontocetes + Mysticeti + Artiodactyla | intensification | 3.64 | 0 | 15.15 |
| | Extant odontocetes | Ancestral odontocetes | intensification | 2.67 | 0.007 | 7.25 |
| CLDN14 | Extant odontocetes | Extant Mysticeti + Artiodactyla | intensification | 1.53 | 0.009 | 6.74 |
| | Extant odontocetes | Ancestral odontocetes | intensification | 1.59 | 0.016 | 5.76 |

**Table S3.** aBSREL results for the genes with evidence of episodic diversifying selection affecting some branches in the tree. B = Optimized branch length. LRT = Likelihood ratio test statistic for selection. Test p-value = p-value corrected for multiple testing.

| Gene | Species | B | LRT | Test p-value | Uncorrected p-value | ω distribution over sites |
|---|---|---|---|---|---|---|
| CDH23 | *Cephalorhynchus commersonii* | 0.0000 | 11.3691 | 0.0152 | 0.0012 | ω1 = 0.0630 (100%) ω2 = 1120 (0.25%) |
| | *Sotalia guianensis* | 0.0000 | 11.2033 | 0.0152 | 0.0013 | ω1 = 0.00 (100%) ω2 = 1350 (0.10%) |
| | *Orcaella heinsohni* | 0.0000 | 10.2908 | 0.0222 | 0.0020 | ω1 = 0.00 (100%) ω2 = 1340 (0.33%) |
| SLC26A5 | *Phocoena sinus* | 0.0000 | 10.8594 | *0.0197* | 0.0015 | ω1 = 0.00 (100%) ω2 = 558 (0.31%) |
| TMC1 | *Neophocaena phocaenoides* | 0 | 329709 | 0 | 0 | ω1 = 0.00 (100%) ω2 = 100000 (0.31%) |

**Table S4.** Positively selected sites on CDH23, coastal dolphins. The column "type of evidence" lists all site tests with significant results - i.e., posterior probability > 0.9 (codeml and FUBAR) and p-value < 0.1 (MEME, FEL, and Contrast-FEL) - that support each PSS.

| Site | Inferred change | Branches with change | Type of evidence |
|---|---|---|---|
| **416** | ATT - GTT Ile - Val | *Neophocaena phocaenoides* *Phocoena phocoena* *Phocoena sinus* | M2, M8, FUBAR, MEME, FEL |
| **1429** | GAC - GTG Asp - Val | *Orcaella heinsohni* | A-model, MEME, FEL, Contrast-Fel, TreeSAAP (Chromatographic index, Hydropathy, Solvent accessible reduction ratio, Surrounding hydrophobicity) |
| **1911** | ACA - GTA Thr - Val | *Sotalia guianensis* | A-model, FUBAR, MEME, FEL, Contrast-FEL, TreeSAAP (Solvent accessible reduction ratio) |
| **2214** | GAG - ACG | *Neophocaena phocaenoides* | FUBAR, MEME, FEL, Contrast-FEL |

| | | | |
|---|---|---|---|
| | Glu - Thr | | |
| | CAG - CAT<br>Gln - His | *Lagenorhynchus obscurus* | |
| 3269 | CAG - CGG<br>Gln - Arg | *Orcaella heinsohni* | A-model, FUBAR, FEL, Contrast-FEL, TreeSAAP (Isoelectric point) |
| **3324** | GCC - ATC<br>Ala - Ile | *Cephalorhynchus commersonii* | FUBAR, MEME, Contrast-FEL, TreeSAAP (Solvent accessible reduction ratio) |

**Table S5.** Positively selected sites on CDH23, riverine dolphins. The column "type of evidence" lists all site tests with significant results - i.e., posterior probability > 0.9 (codeml and FUBAR) and p-value < 0.1 (MEME, FEL, and Contrast-FEL) - that support each PSS.

| Site | Inferred change | Branches with change | Type of evidence |
|---|---|---|---|
| **416** | ATT - GTT<br>Ile - Val | *Lipotes vexillifer*<br>*Neophocaena a. asiaeorientalis* | M2, M8, FUBAR, MEME, FEL |
| **808** | ATT - GTT<br>Ile - Val | *Lipotes vexillifer* | M2, M8, FEL |
| **935** | CCT - AAT<br>Pro - Asn | *Platanista gangetica* | M8, FUBAR, MEME, FEL |
| **1001** | TTC - CAC<br>Phe - His | *Lipotes vexillifer*<br>*Inia geoffrensis*<br>*Orcaella brevirostris*<br>*Sotalia fluviatilis* | FUBAR, FEL, Contrast-FEL |
| | GAT - CAT<br>Asp - His | *Platanista gangetica* | |
| **1370** | GAT - AAT<br>Asp - Asn | *Pontoporia blainvillei* | FUBAR, FEL, Contrast-FEL, TreeSAAP (Isoelectric point) |
| **1446** | ATT - GTT<br>Ile - Val | *Platanista gangetica*<br>*Inia geoffrensis* | M2, M8, FUBAR, MEME, FEL |
| **1450** | CGG - CAG<br>Arg - Gln | *Platanista gangetica* | M8, FUBAR, FEL |
| **1639** | ACG - GTG<br>Thr - Val | *Lipotes vexillifer* | M2, M8, FUBAR, FEL, Contrast-FEL, TreeSAAP (Solvent |

| | | | |
|---|---|---|---|
| | | | accessible reduction ratio) |
| **2288** | AAC - AGC<br>Asn - Ser | *Platanista gangetica*<br>*Inia geoffrensis* | Amodel, FEL, Contrast-FEL |
| **2825** | CCA - GCA<br>Pro - Ala | *Platanista gangetica* | M2, M8, FUBAR, MEME, FEL |
| **3004** | AAG - ATG<br>Lys - Met<br>AAG - CAG<br>Lys - Gln | *Platanista gangetica*<br>*Lipotes vexillifer*<br>*Pontoporia blainvillei* | FUBAR, MEME, FEL, Contrast-FEL, TREESAAP (Chromatographic index, Hydropathy) |

**Table S6.** Positively selected sites on CDH23, oceanic dolphins. The column "type of evidence" lists all site tests with significant results - i.e., posterior probability > 0.9 (codeml and FUBAR) and p-value < 0.1 (MEME, FEL, and Contrast-FEL) - that support each PSS.

| Site | Inferred change | Branches with change | Type of evidence |
|---|---|---|---|
| **410** | GTC - GAC<br>Val - Asp | *Stenella attenuata* | MEME, FEL, TreeSAAP (Polar requirement, Polarity) |
| **782** | AGC - ATC<br>Thr - Ile<br>AGC - CTC<br>Thr - Leu<br>AGC - GTC<br>Thr - Val | *Physeter catodon*<br>*Kogia sima*<br>*Kogia breviceps* | M2, M8, FEl, Contrast-FEL, TreeSAAP (Bulkiness, Chromatographic index, Solvent accessible reduction ratio, Surrounding hydrophobicity) |
| **1450** | CGG - TGG<br>Arg - Trp<br>CGG - TCG<br>Arg - Ser<br>CGG - CAG<br>Arg - Gln | *Physeter catodon*<br>*Kogia sima*<br>*Kogia breviceps*<br>*Hyperoodon ampullatus*<br>*Mesoplodon densirostris* | M8, FUBAR, TreeSAAP (Chromatographic index, Normalized consensus hydrophobicity, Solvent accessible reduction ratio) |
| **1950** | TTG - GTC<br>Leu - Val | *Physeter catodon* | FUBAR, MEME, FEL, Contrast-FEL |
| **2267** | ATT - ACT | *Kogia sima* | M2, M8, FUBAR, FEL |

| | | |
|---|---|---|
| | Ile - Thr | *Kogia breviceps*<br>*Mesoplodon densirostris* |

**Table S7.** Positively selected sites on TMC1, with rows showing coastal, riverine and oceanic dolphins, in this order. The column "type of evidence" lists all site tests with significant results - i.e., posterior probability > 0.9 (codeml and FUBAR) and p-value < 0.1 (MEME, FEL, and Contrast-FEL) - that support each PSS.

| Site | Inferred change | Branches with change | Type of evidence |
|---|---|---|---|
| **344** | AAG - GCA<br>Lys - Ala | *Neophocaena phocaenoides* | M2, M8, FEL, Contrast-FEL, TreeSAAP (Compressibility, Hydropathy) |
| **274** | CTG - ATG<br>Leu - Met | *Platanista gangetica*<br>*Inia geoffrensis*<br>*Lipotes vexillifer*<br>*Pontoporia blainvillei* | A-model, M2, M8, FEL |
| **283** | ATC - GTC<br>Ile - Val | *Platanista gangetica* | FUBAR, FEL |
| **355** | GTA/GCA - GGA<br>Val - Gly | *Pontoporia blainvillei* | FUBAR, FEL |
| **633** | TCG - GGG<br>Ser - Gly | *Neophocaena a. asiaeorientalis* | FUBAR, MEME |
| **129** | AGT - AAT<br>Ser - Asn | *Physeter catodon*<br>*Kogia sima*<br>*Ziphius cavirostris*<br>*Hyperoodon ampullatus*<br>*Mesoplodon densirostris* | M2, FUBAR, Contrast-FEL |
| **151** | | Most species | |
| **274** | GCC - GGC<br>CTG - ATG<br>Leu - Met | *Physeter catodon*<br>*Kogia breviceps* | A-model, M2, M8, FUBAR |

**Table S8.** Positively selected sites on SLC26A5, with rows showing coastal, riverine and oceanic dolphins, in this order. The column "type of evidence" lists all site tests with significant results - i.e., posterior probability > 0.9 (codeml and FUBAR) and p-value < 0.1 (MEME, FEL, and Contrast-FEL) - that support each PSS.

| Site | Inferred change | Branches with change | Type of evidence |
|---|---|---|---|
| **296** | GCG - CAG<br>Ala - Gln | *Phocoena sinus* | FUBAR, MEME, FEL, Contrast-FEL, TreeSAAP (Compressibility(-)) |
| **7** | ATG - GTG (?)<br>Met - Val | *Lipotes vexillifer*<br>*Inia geoffrensis*<br>*Pontoporia blainvillei* | M8, FUBAR |
| **592** | GTC - ACC<br>Val - Thr | *Platanista gangetica* | FUBAR, Contrast-FEL, TreeSAAP (Solvent accessible reduction ratio(-)) |
| **642** | ATC - GTC<br>Ile - Val | *Platanista gangetica*<br>*Pontoporia blainvillei* | FEL, Contrast-FEL |
| **17** | TAT - CTT<br>Tyr - Leu | *Physeter catodon*<br>*Kogia sima*<br>*Kogia breviceps* | FUBAR, MEME |
| **75** | AGA - CAA<br>Lys - Gln | *Kogia breviceps*<br>*Kogia sima* | Isoelectric point(-) |
| **619** | ATA - GTA<br>Ile - Val | *Physeter catodon*<br>*Hyperoodon ampullatus* | FEL, Contrast-FEL |
| **684** | GTG - ATA<br>Val - Ile | *Kogia sima* | MEME, FEL |

**Table S9.** BLAST hits of the genes CDH23, CLDN14, SLC26A4 and TMC1 on the *Sotalia guianensis* and *Sotalia fluviatilis* genomes.

| Query | Length | Location | Length | Start | End | Identity | GAPS |
|---|---|---|---|---|---|---|---|
| ***S.fluviatilis*** | | | | | | | |
| Tursiops_truncatus_TMC1 | 2286 | No hits | | | | | |
| Tursiops_truncatus_SLC26A5 | 2226 | SFLUV_00012249-RA transcript offset:119 AED:0.36 eAED:0.36 | 2153 | 120 | 2000 | 1853/1881 (99%) | 24/1881 (1%) |

| | | | | | | | |
|---|---|---|---|---|---|---|---|
| | | QI:119\|0.82\|0.83\|1\|0.76\|0.88\|18\|0\|677 | | 2044 | 2153 | 107/110 (97%) | 2/110 (2%) |
| Globicephala_melas_SLC26A5 | 2226 | SFLUV_00012249-RA transcript offset:119 AED:0.36 eAED:0.36 QI:119\|0.82\|0.83\|1\|0.76\|0.88\|18\|0\|677 | 2153 | 120 | 2000 | 1852/1881 (98%) | 24/1881 (1%) |
| | | | | 2044 | 2153 | 107/110 (97%) | 2/110 (2%) |
| PREDICTED_Tursiops_truncatus_SMPX_mRNA | 261 | SFLUV_00017804-RA transcript offset:0 AED:0.09 eAED:0.09 QI:0\|0\|0\|0.66\|1\|1\|3\|0\|86 | 261 | 1 | 261 | 259/261 (99%) | 0/261 (0%) |
| PREDICTED_Tursiops_truncatus_CLDN14_mRNA | 711 | SFLUV_00014867-RA transcript offset:0 AED:0.02 eAED:0.02 QI:0\|-1\|0\|1\|-1\|1\|1\|0\|236 | 711 | 1 | 711 | 711/711 (100%) | 0/711 (0%) |
| PREDICTED_Orcinus_orca_CDH23_transcript_variant_X1_mRNA | 10065 | SFLUV_00011738-RA transcript offset:0 AED:0.20 eAED:0.21 QI:0\|0.61\|0.65\|0.96\|0.96\|0.93\|32\|1206\|1789 | 6576 | 2788 | 5370 | 2571/2583 (99%) | 0/2583 (0%) |
| | | | | 1246 | 2788 | 1532/1543 (99%) | 0/1543 (0%) |
| | | | | 736 | 1080 | 344/345 (99%) | 0/345 (0%) |
| | | | | 225 | 570 | 344/346 (99%) | 0/346 (0%) |
| | | | | 575 | 706 | 132/132 (100%) | 0/132 (0%) |
| | | | | 1180 | 1246 | 67/67 (100%) | 0/67 (0%) |
| | | SFLUV_00011742-RA transcript offset:0 AED:0.24 eAED:0.24 QI:0\|0\|0\|1\|1\|1\|11\|0\|564 | 1695 | 68 | 1665 | 1581/1598 (99%) | 0/1598 (0%) |
| | | SFLUV_00011743-RA transcript offset:0 AED:0.08 eAED:0.08 QI:0\|0\|0\|0.66\|0.5\|0.66\|3\|0\|115 | 345 | 3 | 345 | 321/343 (94%) | 18/343 (5%) |
| | | SFLUV_00011741-RA transcript offset:0 AED:0.45 eAED:0.53 QI:0\|0\|0\|0.66\|0.5\|0.66\|3\|0\|154 | 465 | 29 | 291 | 263/263 (100%) | 0/263 (0%) |
| ***S. guianensis*** | | | | | | | |
| Tursiops_truncatus_TMC1 | 2286 | No hits | | | | | |
| Tursiops_truncatus_SLC26A5 | 2226 | SGUI_00011802-RA transcript offset:119 AED:0.36 eAED:0.37 QI:119\|0.87\|0.94\|1\|0.43\|0.64\|17\|0\|645 | 2057 | 120 | 1904 | 1781/1785 (99%) | 0/1785 (0%) |
| | | | | 1948 | 2057 | 107/110 (97%) | 2/110 (2%) |
| | | | | 1901 | 1959 | 58/59 (98%) | 0/59 (0%) |
| PREDICTED_Tursiops_truncatus_SMPX_mRNA | 261 | SGUI_00018365-RA transcript offset:0 AED:0.09 eAED:0.09 QI:0\|0\|0\|0.66\|1\|1\|3\|0\|86 | 261 | 1 | 261 | 259/261 (99%) | 0/261 (0%) |

| | | | | | | | |
|---|---|---|---|---|---|---|---|
| PREDICTED_Tursiops_truncatus_CLDN14_mRNA | 711 | SGUI_00013782-RA transcript offset:0 AED:0.02 eAED:0.02 QI:0\|-1\|0\|1\|-1\|1\|1\|0\|236 | 711 | 1 | 711 | 710/711 (99%) | 0/711 (0%) |
| PREDICTED_Orcinus_orca_CDH23_transcript_variant_X1_mRNA | 10065 | SGUI_00010396-RA transcript offset:0 AED:0.21 eAED:0.22 QI:0\|0.61\|0.65\|0.96\|0.96\|0.93\|32\|713\|1789 | 6083 | 2788 | 5370 | 2571/2583 (99%) | 0/2583 (0%) |
| | | | | 1246 | 2788 | 1532/1543 (99%) | 0/1543 (0%) |
| | | | | 736 | 1080 | 345/345 (100%) | 0/345 (0%) |
| | | | | 225 | 570 | 344/346 (99%) | 0/346 (0%) |
| | | | | 575 | 706 | 132/132 (100%) | 0/132 (0%) |
| | | | | 1180 | 1246 | 67/67 (100%) | 0/67 (0%) |
| | | SGUI_00010400-RA transcript offset:0 AED:0.15 eAED:0.18 QI:0\|0\|0\|1\|0.91\|1\|13\|0\|699 | 2100 | 68 | 1926 | 1842/1859 (99%) | 0/1859 (0%) |
| | | SGUI_00010401-RA transcript offset:0 AED:0.35 eAED:0.35 QI:0\|0\|0\|0.25\|1\|1\|8\|0\|343 | 1029 | 1 | 1029 | 1000/1044 (96%) | 39/1044 (4%) |
| | | SGUI_00010398-RA transcript offset:0 AED:0.35 eAED:0.35 QI:0\|0\|0\|0.16\|0.54\|0.58\|12\|0\|598 | 1797 | 1130 | 1620 | 489/491 (99%) | 0/491 (0%) |
| | | | | 379 | 852 | 471/492 (96%) | 18/492 (4%) |
| | | | | 1 | 346 | 342/346 (99%) | 0/346 (0%) |
| | | | | 926 | 1081 | 156/156 (100%) | 0/156 (0%) |

**Table S10.** BLAST hits of the genes CDH23, CLDN14, SLC26A4 and TMC1 on the *Sotalia guianensis* and *Sotalia fluviatilis* genomes.

| Query | Location | Length | Start | End | Identity | GAPS |
|---|---|---|---|---|---|---|
| *S.fluviatilis* | | | | | | |

| | | | | | | |
|---|---|---|---|---|---|---|
| Globicephala melas_TMC1 | scaffold_2034_uid_1558006649 | 241 | 118294 | 118534 | 239/241 (99%) | 0/241 (0%) |
| | | 197 | 93022 | 93218 | 197/197 (100%) | 0/197 (0%) |
| | | 180 | 95438 | 95617 | 180/180 (100%) | 0/180 (0%) |
| | | 183 | 34907 | 35089 | 181/183 (99%) | 0/183 (0%) |
| | | 169 | 95741 | 95909 | 168/169 (99%) | 1/169 (1%) |
| | | 146 | 92261 | 92406 | 146/146 (100%) | 0/146 (0%) |
| | | 144 | 87692 | 87835 | 144/144 (100%) | 0/144 (0%) |
| | | 132 | 102745 | 102876 | 132/132 (100%) | 0/132 (0%) |
| | | 130 | 39634 | 39763 | 130/130 (100%) | 0/130 (0%) |
| | | 131 | 123279 | 123409 | 130/131 (99%) | 0/131 (0%) |
| | | 115 | 72249 | 72363 | 113/115 (98%) | 2/115 (2%) |
| | | 102 | 73463 | 73564 | 102/102 (100%) | 0/102 (0%) |
| | | 96 | 58089 | 58184 | 95/96 (99%) | 0/96 (0%) |
| | | 90 | 59871 | 59960 | 89/90 (99%) | 0/90 (0%) |
| | | 79 | 127402 | 127480 | 79/79 (100%) | 0/79 (0%) |
| | | 72 | 113352 | 113423 | 70/72 (97%) | 0/72 (0%) |
| PREDICTED_Orcinus_orca_CDH23_transcript_variant_X1_mRNA | scaffold_3512_uid_1558006649 (plus minus) | 468 | 456840 | 456374 | 463/468 (99%) | 1/468 (0%) |
| | | 391 | 517132 | 516742 | 389/391 (99%) | 0/391 (0%) |
| | | 329 | 437309 | 436981 | 326/329 (99%) | 0/329 (0%) |
| | | 258 | 441424 | 441167 | 257/258 (99%) | 0/258 (0%) |
| | | 256 | 443263 | 443008 | 254/256 (99%) | 0/256 (0%) |
| | | 243 | 564496 | 564254 | 241/243 (99%) | 0/243 (0%) |
| | | 231 | 507850 | 507620 | 230/231 (99%) | 0/231 (0%) |
| | | 227 | 451218 | 450992 | 225/227 (99%) | 0/227 (0%) |
| | | 224 | 470446 | 470223 | 222/224 (99%) | 0/224 (0%) |
| | | 223 | 539677 | 539455 | 221/223 (99%) | 0/223 (0%) |
| | | 214 | 446926 | 446713 | 213/214 (99%) | 0/214 (0%) |
| | | 221 | 463055 | 462836 | 218/221 (99%) | 1/221 (0%) |

| | | | | | | |
|---|---|---|---|---|---|---|
| | | 206 | 458554 | 458349 | 203/206 (99%) | 0/206 (0%) |
| | | 203 | 708299 | 708098 | 201/203 (99%) | 1/203 (0%) |
| | | 194 | 444713 | 444520 | 194/194 (100%) | 0/194 (0%) |
| | | 191 | 656837 | 656647 | 191/191 (100%) | 0/191 (0%) |
| | | 192 | 542694 | 542503 | 191/192 (99%) | 0/192 (0%) |
| | | 188 | 469156 | 468969 | 187/188 (99%) | 0/188 (0%) |
| | | 181 | 447255 | 447075 | 179/181 (99%) | 0/181 (0%) |
| | | 173 | 450449 | 450277 | 173/173 (100%) | 0/173 (0%) |
| | | 172 | 442935 | 442764 | 171/172 (99%) | 0/172 (0%) |
| | | 160 | 630979 | 630820 | 160/160 (100%) | 0/160 (0%) |
| | | 156 | 511652 | 511497 | 155/156 (99%) | 0/156 (0%) |
| | | 157 | 537708 | 537552 | 155/157 (99%) | 0/157 (0%) |
| | | 150 | 533608 | 533459 | 150/150 (100%) | 0/150 (0%) |
| | | 151 | 631619 | 631469 | 150/151 (99%) | 0/151 (0%) |
| | | 149 | 542221 | 542073 | 148/149 (99%) | 0/149 (0%) |
| | | 147 | 450058 | 449913 | 146/147 (99%) | 1/147 (1%) |
| | | 150 | 523196 | 523043 | 150/154 (97%) | 0/154 (0%) |
| | | 159 | 518824 | 518668 | 153/159 (96%) | 3/159 (2%) |
| | | 138 | 463734 | 463597 | 136/138 (99%) | 0/138 (0%) |
| | | 131 | 443493 | 443363 | 131/131 (100%) | 0/131 (0%) |
| | | 138 | 509130 | 508993 | 136/138 (99%) | 1/138 (1%) |
| | | 134 | 510178 | 510045 | 133/134 (99%) | 0/134 (0%) |
| | | 131 | 704490 | 704360 | 131/131 (100%) | 0/131 (0%) |
| | | 130 | 561159 | 561030 | 130/130 (100%) | 0/130 (0%) |
| | | 128 | 459391 | 459264 | 128/128 (100%) | 0/128 (0%) |
| | | 129 | 438829 | 438701 | 128/129 (99%) | 0/129 (0%) |
| | | 132 | 439141 | 439010 | 130/132 (98%) | 0/132 (0%) |
| | | 124 | 469932 | 469809 | 124/124 (100%) | 0/124 (0%) |

| | | | | | | |
|---|---|---|---|---|---|---|
| | | 122 | 440302 | 440181 | 122/122 (100%) | 0/122 (0%) |
| | | 121 | 452578 | 452458 | 121/121 (100%) | 0/121 (0%) |
| | | 123 | 448924 | 448802 | 122/123 (99%) | 0/123 (0%) |
| | | 118 | 444352 | 444236 | 117/118 (99%) | 1/118 (1%) |
| | | 117 | 658633 | 658517 | 116/117 (99%) | 0/117 (0%) |
| | | 123 | 554650 | 554528 | 119/123 (97%) | 0/123 (0%) |
| | | 107 | 438275 | 438169 | 107/107 (100%) | 0/107 (0%) |
| | | 114 | 462291 | 462179 | 112/114 (98%) | 1/114 (1%) |
| | | 117 | 548893 | 548777 | 114/117 (97%) | 1/117 (1%) |
| | | 111 | 562865 | 562755 | 109/111 (98%) | 0/111 (0%) |
| | | 103 | 515902 | 515800 | 102/103 (99%) | 0/103 (0%) |
| | | 105 | 460781 | 460677 | 103/105 (98%) | 0/105 (0%) |
| | | 101 | 440820 | 440720 | 100/101 (99%) | 0/101 (0%) |
| | | 82 | 440100 | 440019 | 82/82 (100%) | 0/82 (0%) |
| | | 84 | 695836 | 695753 | 82/84 (98%) | 0/84 (0%) |
| | | 66 | 439687 | 439622 | 66/66 (100%) | 0/66 (0%) |
| | | 66 | 524225 | 524160 | 66/66 (100%) | 0/66 (0%) |
| | | 68 | 594462 | 594395 | 67/68 (99%) | 0/68 (0%) |
| | scaffold_1613_uid_1558006649 (plus plus) | 144 | 2047449 | 2047592 | 144/144 (100%) | 0/144 (0%) |
| | | 95 | 2048431 | 2048525 | 94/95 (99%) | 0/95 (0%) |
| | | 81 | 1988217 | 1988297 | 81/81 (100%) | 0/81 (0%) |
| | | 65 | 1981791 | 1981855 | 65/65 (100%) | 0/65 (0%) |
| Globicephala_melas_SLC26A5 | scaffold_111_uid_1558006649 | 202 | 306953 | 307154 | 201/202 (99%) | 0/202 (0%) |
| | | 186 | 309034 | 309219 | 186/186 (100%) | 0/186 (0%) |
| | | 169 | 286092 | 286260 | 169/169 (100%) | 0/169 (0%) |
| | | 175 | 286809 | 286981 | 173/175 (99%) | 2/175 (1%) |
| | | 152 | 279073 | 279224 | 152/152 (100%) | 0/152 (0%) |
| | | 155 | 288925 | 289079 | 154/155 (99%) | 0/155 (0%) |

| | | | | | | |
|---|---|---|---|---|---|---|
| | | 150 | 296495 | 296644 | 149/150 (99%) | 0/150 (0%) |
| | | 145 | 281185 | 281329 | 144/145 (99%) | 0/145 (0%) |
| | | 122 | 284887 | 285007 | 120/122 (98%) | 2/122 (2%) |
| | | 116 | 297694 | 297809 | 115/116 (99%) | 0/116 (0%) |
| | | 113 | 305965 | 306077 | 113/113 (100%) | 0/113 (0%) |
| | | 108 | 299668 | 299775 | 107/108 (99%) | 0/108 (0%) |
| | | 99 | 299358 | 299456 | 99/99 (100%) | 0/99 (0%) |
| | | 95 | 305147 | 305241 | 94/95 (99%) | 0/95 (0%) |
| | | 85 | 291767 | 291851 | 85/85 (100%) | 0/85 (0%) |
| | | 82 | 298600 | 298681 | 82/82 (100%) | 0/82 (0%) |
| | | 74 | 303707 | 303780 | 74/74 (100%) | 0/74 (0%) |
| *S.guianensis* | | | | | | |
| Globicephala melas_TMC1 | scaffold_4522_uid_1568486962 | 241 | 35649 | 35889 | 239/241 (99%) | 0/241 (0%) |
| | | 197 | 10393 | 10589 | 197/197 (100%) | 0/197 (0%) |
| | | 180 | 12808 | 12987 | 180/180 (100%) | 0/180 (0%) |
| | | 169 | 13111 | 13279 | 168/169 (99%) | 1/169 (1%) |
| | | 146 | 9632 | 9777 | 146/146 (100%) | 0/146 (0%) |
| | | 144 | 5063 | 5206 | 144/144 (100%) | 0/144 (0%) |
| | | 132 | 20118 | 20249 | 132/132 (100%) | 0/132 (0%) |
| | | 131 | 40635 | 40765 | 130/131 (99%) | 0/131 (0%) |
| | | 79 | 44753 | 44831 | 79/79 (100%) | 0/79 (0%) |
| | | 72 | 30707 | 30778 | 70/72 (97%) | 0/72 (0%) |
| | scaffold_1193_uid_1568486962 | 183 | 1076010 | 1076192 | 181/183 (99%) | 0/183 (0%) |
| | | 130 | 1080742 | 1080871 | 129/130 (99%) | 0/130 (0%) |
| | | 115 | 1113319 | 1113433 | 113/115 (98%) | 2/115 (2%) |
| | | 102 | 1114533 | 1114634 | 102/102 (100%) | 0/102 (0%) |
| | | 96 | 1099173 | 1099268 | 95/96 (99%) | 0/96 (0%) |
| | | 90 | 1100955 | 1101044 | 88/90 (98%) | 0/90 (0%) |

| | | | | | | |
|---|---|---|---|---|---|---|
| Globicephala_melas_SLC26A5 | scaffold_1486_uid_1568486962 | 202 | 339233 | 339032 | 201/202 (99%) | 0/202 (0%) |
| | | 186 | 337152 | 336967 | 185/186 (99%) | 0/186 (0%) |
| | | 169 | 360064 | 359896 | 169/169 (100%) | 0/169 (0%) |
| | | 175 | 359347 | 359175 | 173/175 (99%) | 2/175 (1%) |
| | | 155 | 357231 | 357077 | 154/155 (99%) | 0/155 (0%) |
| | | 152 | 367085 | 366934 | 152/152 (100%) | 0/152 (0%) |
| | | 149 | 349691 | 349542 | 149/150 (99%) | 0/150 (0%) |
| | | 142 | 364970 | 364829 | 142/142 (100%) | 0/142 (0%) |
| | | 115 | 361264 | 361150 | 115/115 (100%) | 0/115 (0%) |
| | | 113 | 340221 | 340109 | 113/113 (100%) | 0/113 (0%) |
| | | 116 | 348492 | 348377 | 115/116 (99%) | 0/116 (0%) |
| | | 108 | 346518 | 346411 | 107/108 (99%) | 0/108 (0%) |
| | | 99 | 346828 | 346730 | 99/99 (100%) | 0/99 (0%) |
| | | 95 | 341039 | 340945 | 94/95 (99%) | 0/95 (0%) |
| | | 85 | 354389 | 354305 | 85/85 (100%) | 0/85 (0%) |
| | | 82 | 347586 | 347505 | 82/82 (100%) | 0/82 (0%) |
| | | 74 | 342479 | 342406 | 74/74 (100%) | 0/74 (0%) |
| PREDICTED_Orcinus_orca_CDH23_transcript_variant_X1_mRNA | scaffold_344_uid_1568486962 | 468 | 456648 | 456182 | 463/468 (99%) | 1/468 (0%) |
| | | 391 | 517027 | 516637 | 389/391 (99%) | 0/391 (0%) |
| | | 329 | 437183 | 436855 | 326/329 (99%) | 0/329 (0%) |
| | | 258 | 441298 | 441041 | 257/258 (99%) | 0/258 (0%) |
| | | 256 | 443137 | 442882 | 254/256 (99%) | 0/256 (0%) |
| | | 243 | 564395 | 564153 | 240/243 (99%) | 0/243 (0%) |
| | | 231 | 507745 | 507515 | 230/231 (99%) | 0/231 (0%) |
| | | 227 | 451091 | 450865 | 225/227 (99%) | 0/227 (0%) |
| | | 224 | 470248 | 470025 | 222/224 (99%) | 0/224 (0%) |
| | | 223 | 539570 | 539348 | 221/223 (99%) | 0/223 (0%) |
| | | 214 | 446799 | 446586 | 213/214 (99%) | 0/214 (0%) |

| | | | | | | |
|---|---|---|---|---|---|---|
| | | 221 | 462861 | 462642 | 218/221 (99%) | 1/221 (0%) |
| | | 206 | 458361 | 458156 | 203/206 (99%) | 0/206 (0%) |
| | | 194 | 444586 | 444393 | 194/194 (100%) | 0/194 (0%) |
| | | 203 | 707797 | 707596 | 200/203 (99%) | 1/203 (0%) |
| | | 191 | 656370 | 656180 | 191/191 (100%) | 0/191 (0%) |
| | | 192 | 542587 | 542396 | 191/192 (99%) | 0/192 (0%) |
| | | 188 | 468958 | 468771 | 187/188 (99%) | 0/188 (0%) |
| | | 181 | 447128 | 446948 | 179/181 (99%) | 0/181 (0%) |
| | | 173 | 450322 | 450150 | 173/173 (100%) | 0/173 (0%) |
| | | 172 | 442809 | 442638 | 171/172 (99%) | 0/172 (0%) |
| | | 160 | 630476 | 630317 | 160/160 (100%) | 0/160 (0%) |
| | | 156 | 511547 | 511392 | 156/156 (100%) | 0/156 (0%) |
| | | 157 | 537601 | 537445 | 155/157 (99%) | 0/157 (0%) |
| | | 150 | 533502 | 533353 | 150/150 (100%) | 0/150 (0%) |
| | | 151 | 631116 | 630966 | 150/151 (99%) | 0/151 (0%) |
| | | 149 | 542114 | 541966 | 148/149 (99%) | 0/149 (0%) |
| | | 147 | 449931 | 449786 | 146/147 (99%) | 1/147 (1%) |
| | | 154 | 523091 | 522938 | 150/154 (97%) | 0/154 (0%) |
| | | 159 | 518719 | 518563 | 153/159 (96%) | 3/159 (2%) |
| | | 138 | 463540 | 463403 | 137/138 (99%) | 0/138 (0%) |
| | | 131 | 443366 | 443236 | 131/131 (100%) | 0/131 (0%) |
| | | 138 | 509025 | 508888 | 136/138 (99%) | 1/138 (1%) |
| | | 134 | 510073 | 509940 | 133/134 (99%) | 0/134 (0%) |
| | | 131 | 703990 | 703860 | 131/131 (100%) | 0/131 (0%) |
| | | 130 | 561058 | 560929 | 130/130 (100%) | 0/130 (0%) |
| | | 128 | 459198 | 459071 | 128/128 (100%) | 0/128 (0%) |
| | | 129 | 438703 | 438575 | 128/129 (99%) | 0/129 (0%) |
| | | 132 | 439015 | 438884 | 130/132 (98%) | 0/132 (0%) |

| | | | | | | |
|---|---|---|---|---|---|---|
| | | 124 | 469734 | 469611 | 124/124 (100%) | 0/124 (0%) |
| | | 122 | 440176 | 440055 | 122/122 (100%) | 0/122 (0%) |
| | | 121 | 452451 | 452331 | 121/121 (100%) | 0/121 (0%) |
| | | 123 | 448797 | 448675 | 122/123 (99%) | 0/123 (0%) |
| | | 120 | 550313 | 550194 | 120/120 (100%) | 0/120 (0%) |
| | | 119 | 535405 | 535287 | 119/119 (100%) | 0/119 (0%) |
| | | 118 | 444225 | 444109 | 117/118 (99%) | 1/118 (1%) |
| | | 117 | 658166 | 658050 | 116/117 (99%) | 0/117 (0%) |
| | | 123 | 554548 | 554426 | 119/123 (97%) | 0/123 (0%) |
| | | 117 | 548790 | 548674 | 115/117 (98%) | 1/117 (1%) |
| | | 107 | 438149 | 438043 | 107/107 (100%) | 0/107 (0%) |
| | | 114 | 462097 | 461985 | 112/114 (98%) | 1/114 (1%) |
| | | 111 | 562764 | 562654 | 109/111 (98%) | 0/111 (0%) |
| | | 103 | 515797 | 515695 | 102/103 (99%) | 0/103 (0%) |
| | | 105 | 460587 | 460483 | 103/105 (98%) | 0/105 (0%) |
| | | 101 | 440694 | 440594 | 100/101 (99%) | 0/101 (0%) |
| | | 82 | 439974 | 439893 | 82/82 (100%) | 0/82 (0%) |
| | | 84 | 695336 | 695253 | 82/84 (98%) | 0/84 (0%) |
| | | 66 | 439561 | 439496 | 66/66 (100%) | 0/66 (0%) |
| | | 66 | 524120 | 524055 | 66/66 (100%) | 0/66 (0%) |
| | | 68 | 593933 | 593866 | 67/68 (99%) | 0/68 (0%) |
| | scaffold_652_uid_1568486962 | 144 | 570282 | 570425 | 144/144 (100%) | 0/144 (0%) |
| | | 95 | 571264 | 571358 | 94/95 (99%) | 0/95 (0%) |
| | | 81 | 511060 | 511140 | 81/81 (100%) | 0/81 (0%) |
| | | 65 | 504631 | 504695 | 65/65 (100%) | 0/65 (0%) |

**Table S11.** Contrast-FEL results for differential selection among river and marine dolphins. The column R x M depicts the sites that showed significantly different ω ratios among these two groups. ω riverine, ω marine and ω background represent the ω values for these sites for each dolphin group and for the background lineages, respectively.

| **Gene** | **Total sites** | **Sites with different dN/dS ratios** | | | | **p-value** |
|---|---|---|---|---|---|---|
| | | R x M | ω riverine | ω marine | ω background | |
| CDH23 | 3365 | 15 | 0.176 | 0.168 | 0.104 | 0.1 |
| SLC26A5 | 747 | 2 | 0.132 | 0.144 | 0.157 | 0.1 |
| TMC1 | 766 | 3 | 0.382 | 0.124 | 0.125 | 0.1 |
| CLDN14 | 243 | 1 | 0.0273 | 0.0156 | 0.0919 | 0.1 |

**Table S12.** Retrieved sequences for 42 cetacean species used in this study, including 37 toothed whales and five baleen whales, with the Genbank accession numbers for each sequence.

https://docs.google.com/spreadsheets/d/1d410SD-iS0vJQDKpPhGh_Uzww2UDAOuizsshoxsorVU/edit?usp=sharing

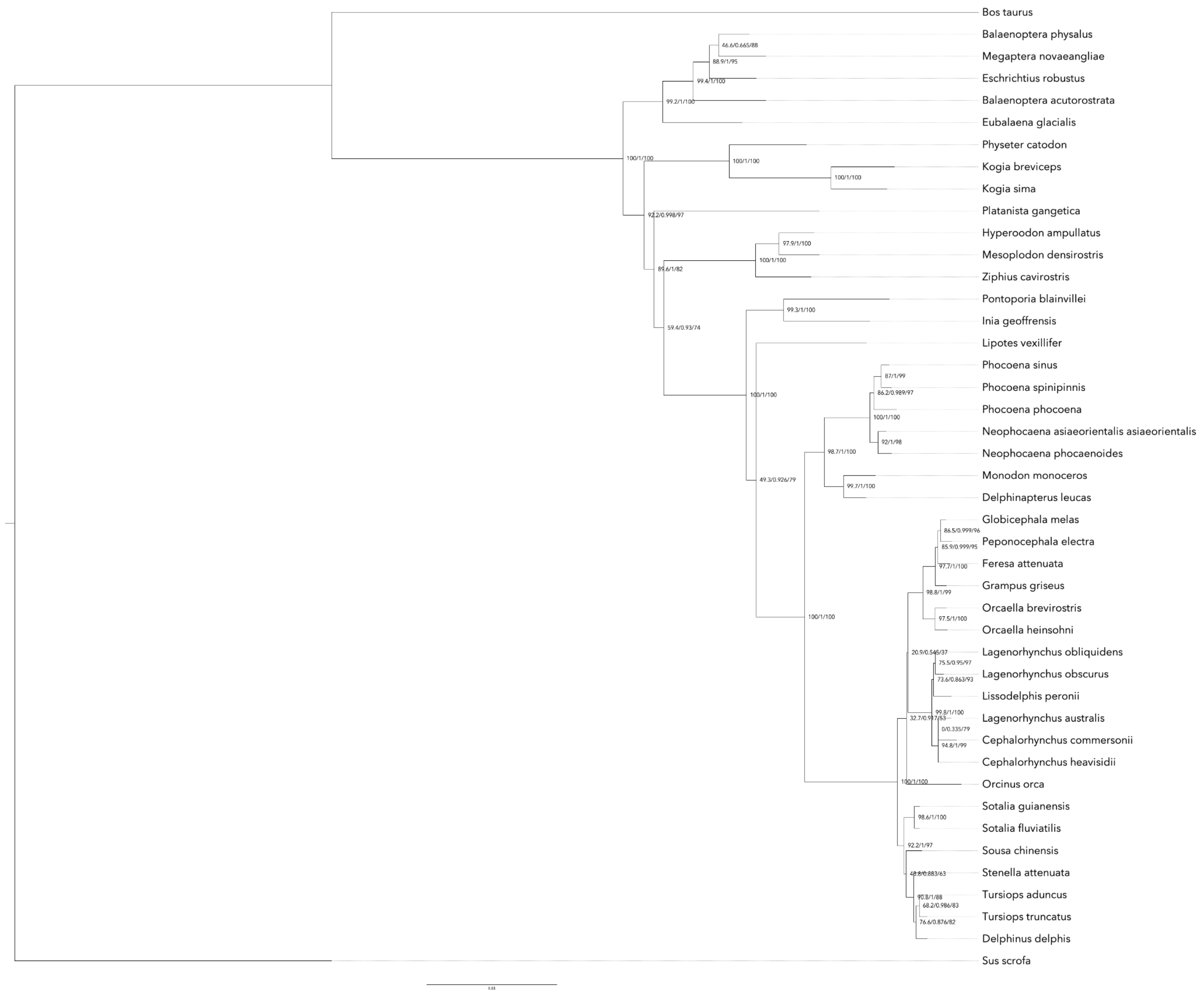

**Figure S1.** Nucleotide tree, CDH23

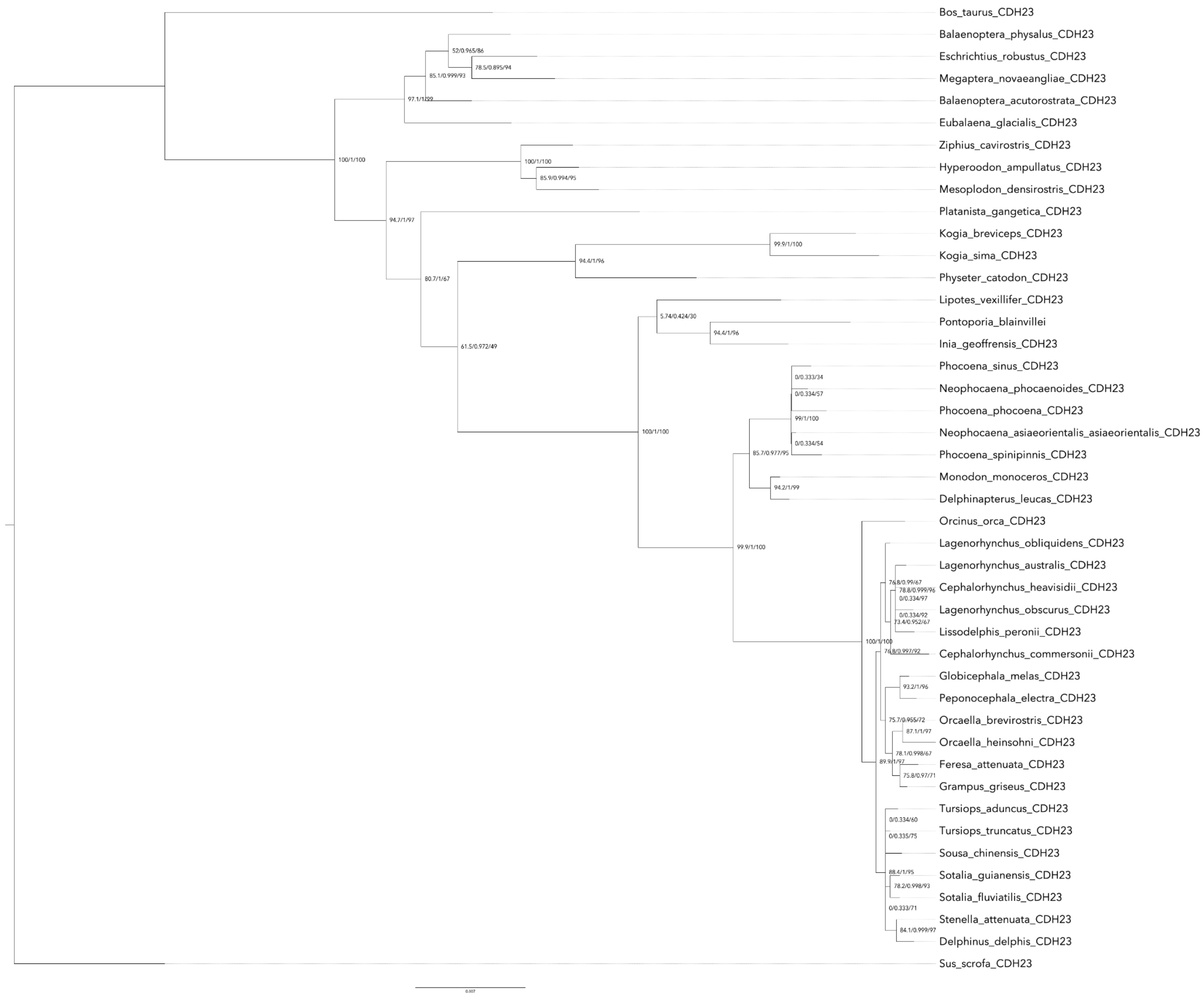

**Figure S2.** Protein tree, CDH23

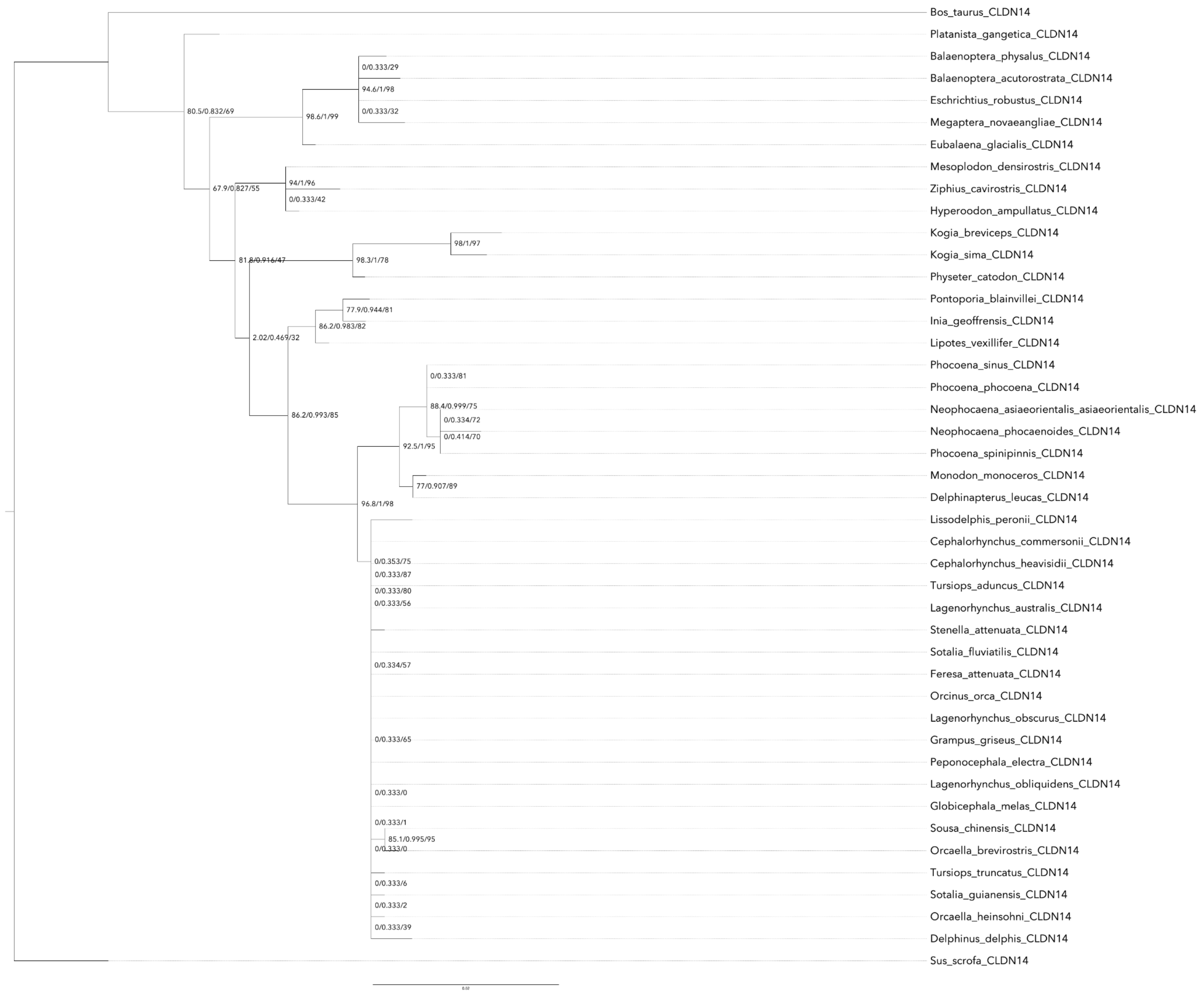

**Figure S3.** Nucleotide tree, CLDN14

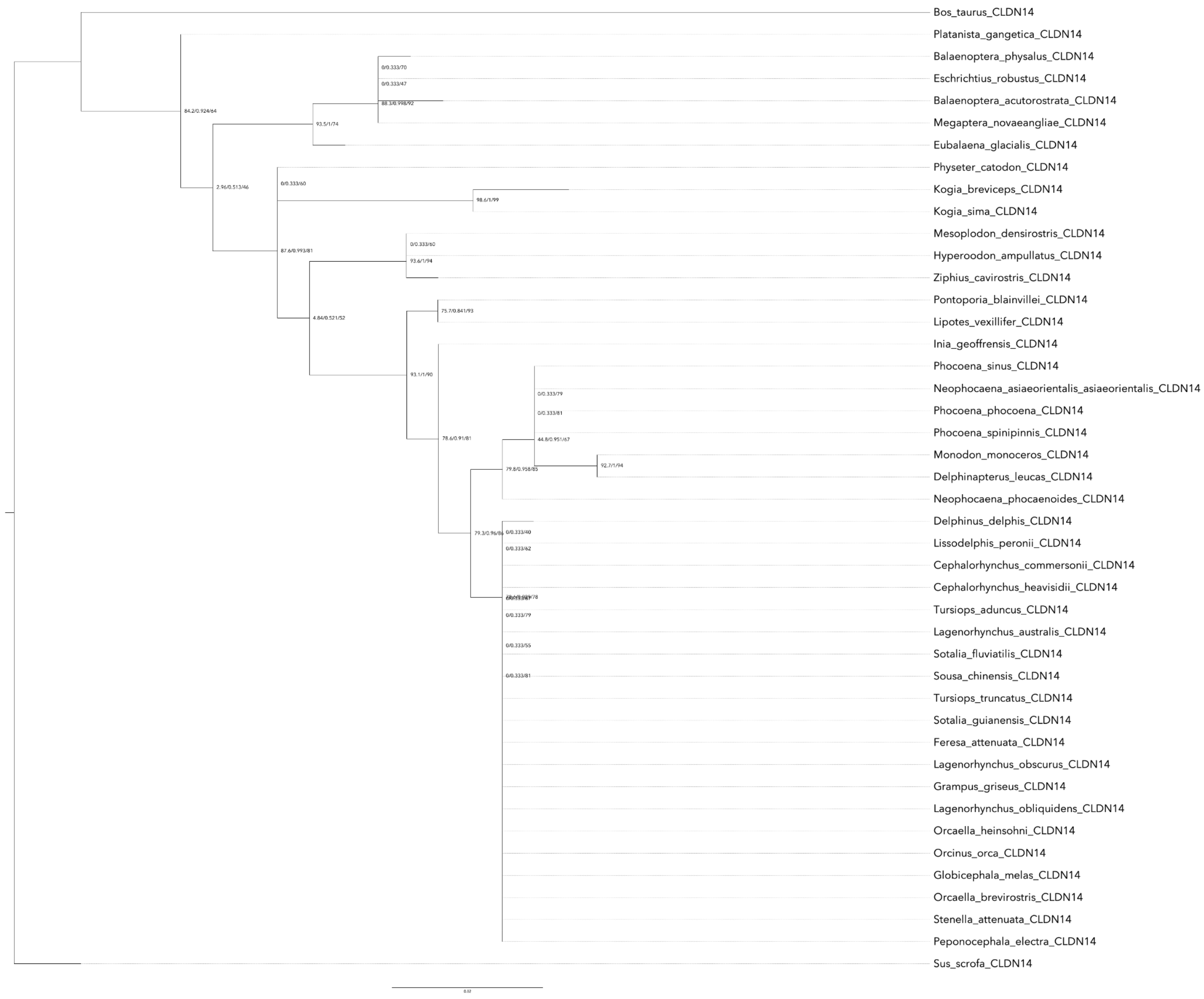

**Figure S4.** Protein tree, CLDN14

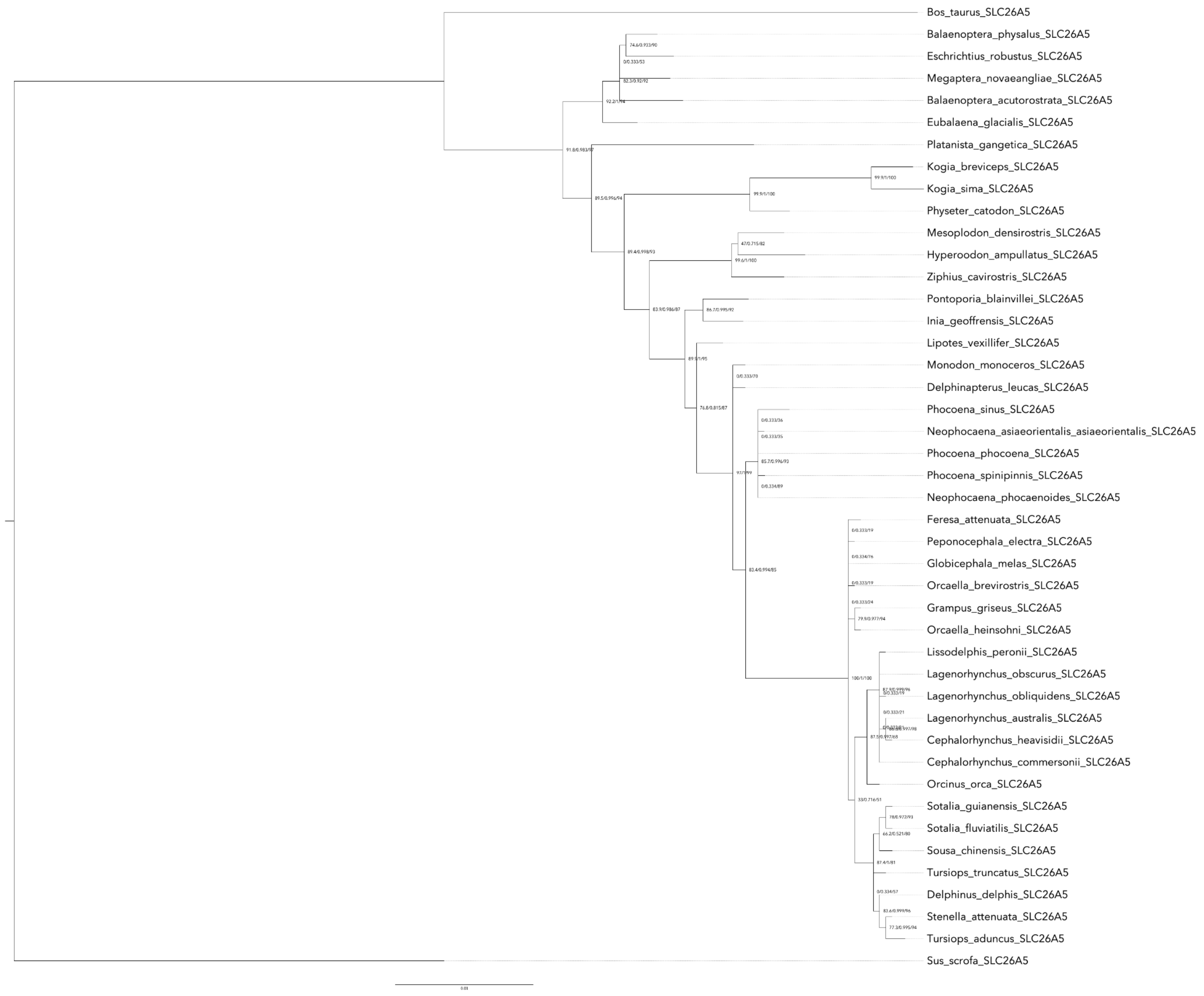

**Figure S5.** Nucleotide tree, SLC26A5

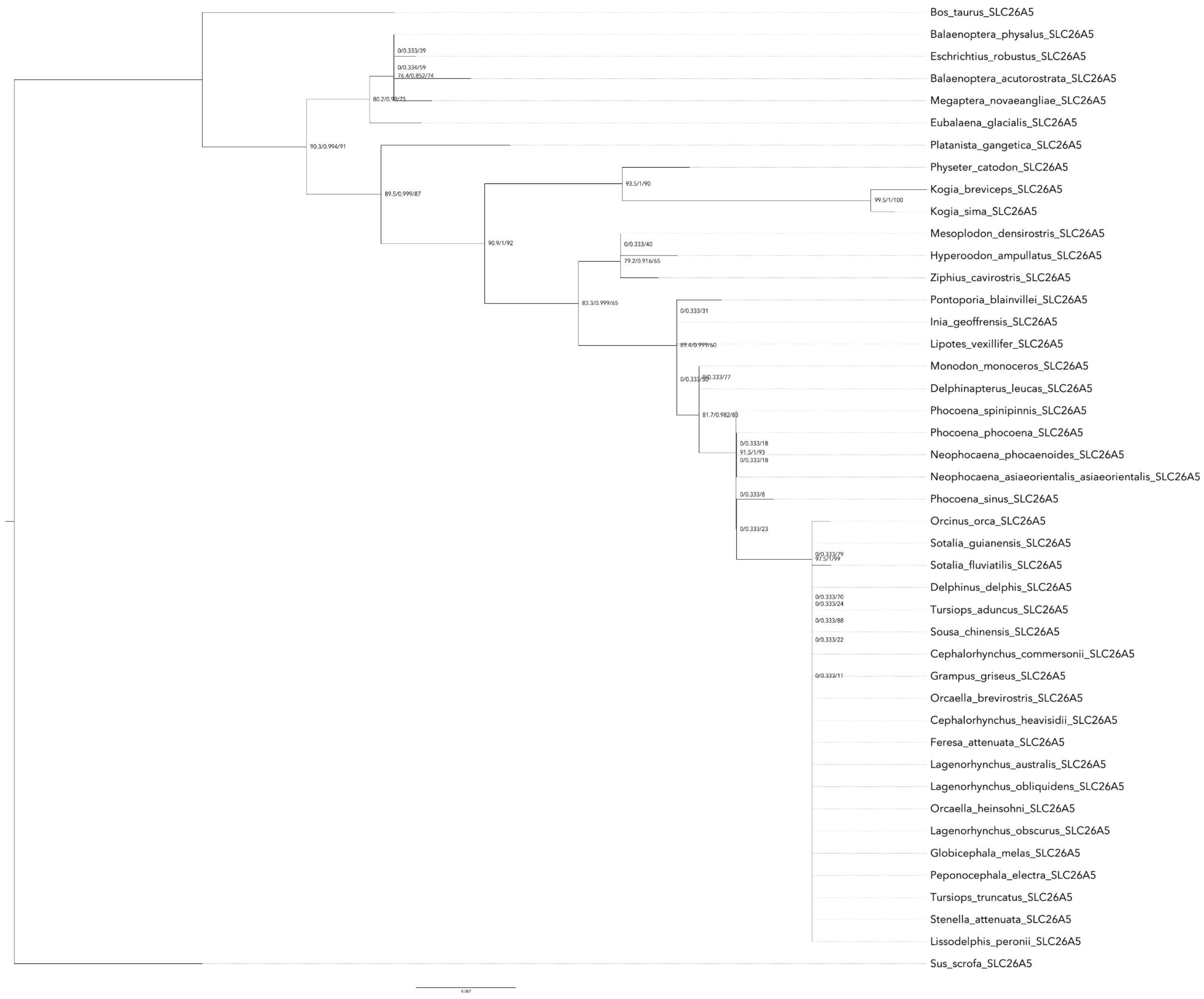

**Figure S6.** Protein tree, SLC26A5

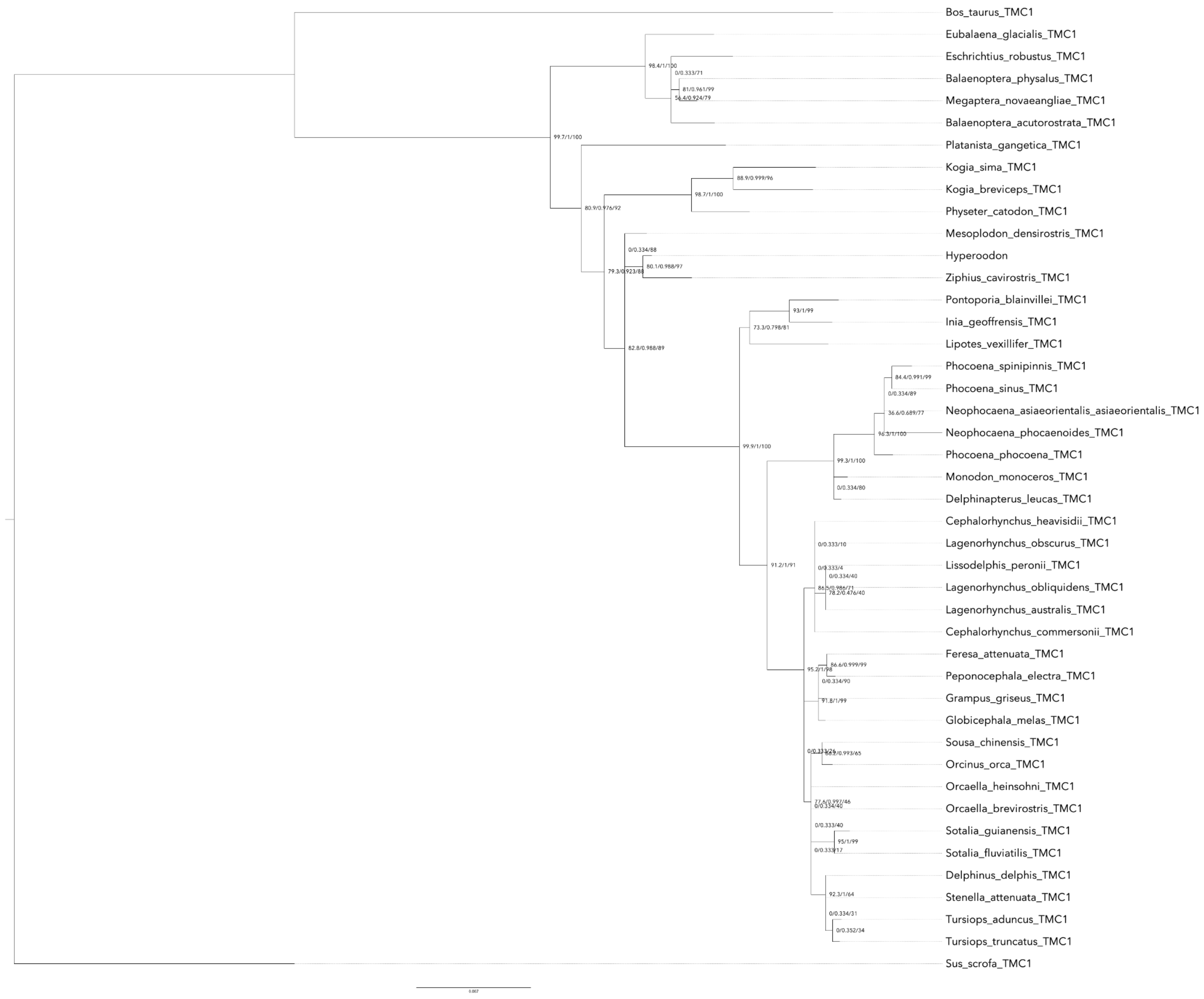

**Figure S7.** Nucleotide tree, TMC1

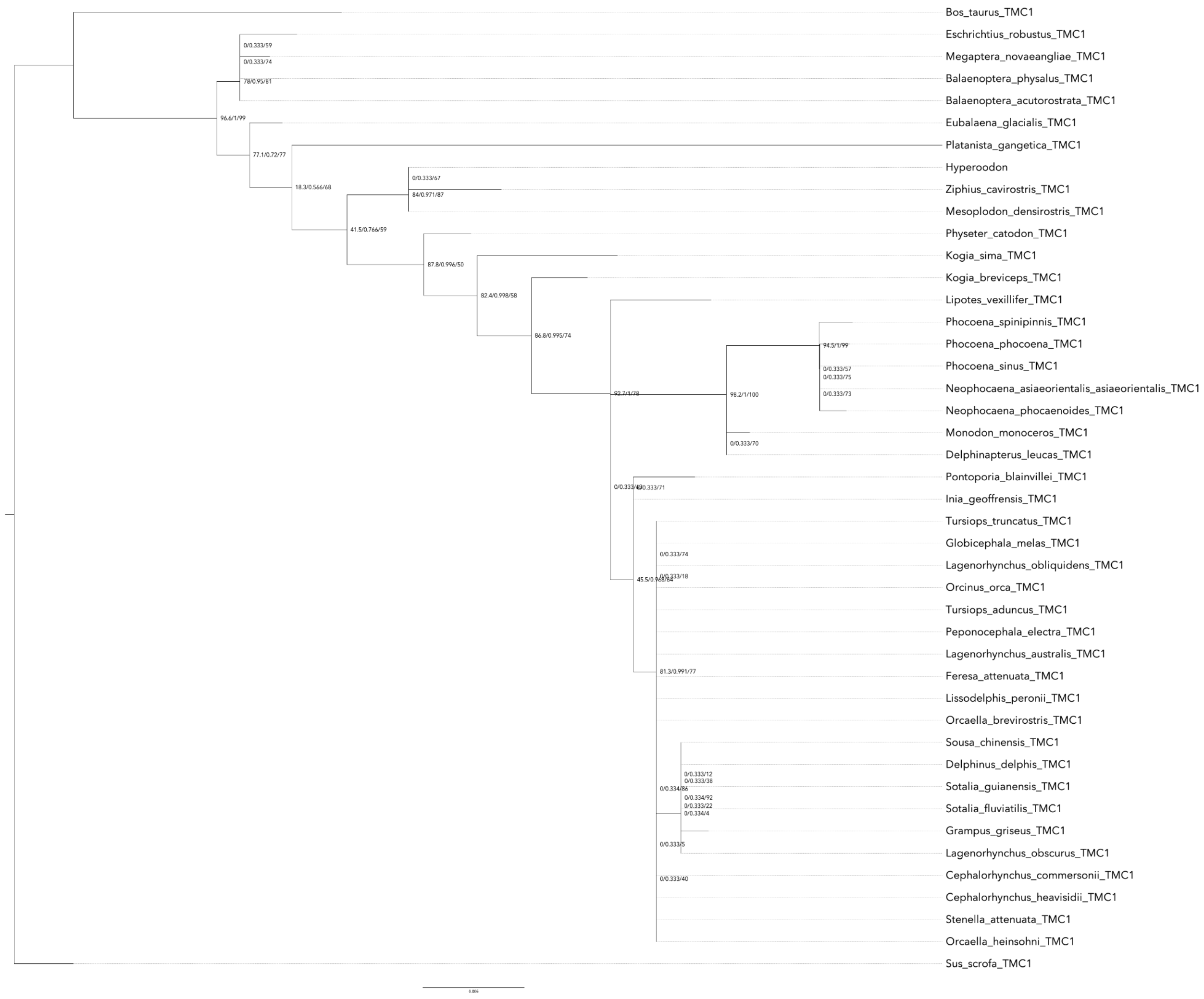

**Figure S7.** Protein tree, TMC1

1. Jensen FH, Johnson M, Ladegaard M, Wisniewska DM, Madsen PT. Narrow Acoustic Field of View Drives Frequency Scaling in Toothed Whale Biosonar. Curr Biol. 2018;28:3878–85.e3.
